# Mean daily temperatures can predict the thermal limits of malaria transmission better than rate summation

**DOI:** 10.1101/2024.09.20.614098

**Authors:** Marta S. Shocket, Joey R. Bernhardt, Kerri L. Miazgowicz, Alyzeh Orakzai, Van M. Savage, Richard J. Hall, Sadie J. Ryan, Courtney C. Murdock

## Abstract

Temperature shapes the distribution, seasonality, and magnitude of mosquito-borne disease outbreaks. Mechanistic models predicting transmission often use mosquito and pathogen thermal responses from constant temperature experiments. However, mosquitoes live in fluctuating environments. Rate summation (nonlinear averaging) is a common approach to infer performance in fluctuating environments, but its accuracy is rarely validated. We measured three mosquito traits that impact transmission (bite rate, survival, fecundity) in a malaria mosquito (*Anopheles stephensi*) across temperature gradients with three diurnal temperature ranges (0, 9 and 12°C). We compared thermal suitability models with temperature-trait relationships observed under constant temperatures, fluctuating temperatures, and those predicted by rate summation. We mapped results across *An. stephenesi*’s native Asian and invasive African ranges. We found: 1) daily temperature fluctuation significantly altered trait thermal responses; 2) rate summation partially captured decreases in performance near thermal optima, but also incorrectly predicted increases near thermal limits; and 3) while thermal suitability characterized across constant temperatures did not perfectly capture suitability in fluctuating environments, it was more accurate for estimating and mapping thermal limits than predictions from rate summation. Our study provides insight into methods for predicting mosquito-borne disease risk and emphasizes the need to improve understanding of organismal performance under fluctuating conditions.

## Introduction

Malaria remains one of the biggest global public health burdens, despite substantial control efforts. In 2022 alone, there were 249 million cases and 580,000 deaths worldwide, mostly of children under five years of age (76% of deaths) and occurring in Africa (94% of cases)^1^. Further, global climate change and land use change are altering the environments where malaria is transmitted, shifting the times of year and geographic regions that are environmentally suitable for malaria transmission^2–7^. Over the past 20 years, we have gained substantial mechanistic insight into how key abiotic environmental variables–including temperature–shape malaria risk^2,6,8–12^. Because mosquitoes are ectothermic, temperature has strong effects on the vital rates of both the mosquito and the parasite. These effects shape mosquito population dynamics, the ability of the mosquito to become infected and transmit, and the parasite development rate, all of which in turn influence malaria transmission dynamics. Thus, a mechanistic determination of how temperature will alter the distribution and abundance of mosquito vectors, as well as people’s potential exposure to malaria-infectious mosquitoes, will be critical for accurately anticipating how the environmental suitability for malaria transmission will respond to current and future global change.

Previous empirical work has focused on characterizing the effects of temperature on mosquito and parasite traits that are relevant for transmission across a diversity of mosquito-borne disease systems^4^. In general, temperature-trait relationships have a kinetic profile akin to an enzymatic reaction^13,14^. Performance is constrained by a lower and upper temperature threshold (*T_min_* and *T_max_*, respectively) and gradually increases with temperature to an optimal value (*T_opt_*) as enzymatic and biochemical processes become more efficient. Performance then declines as temperatures warm away from the *T_opt_*, presumably because enzymatic reactions become less efficient as protein stability declines, followed by performance failure or organism death as temperatures approach the *T_max_*^15–18^. Collectively, these responses give us temperature-trait relationships known as thermal performance curves (TPCs) that have been used extensively across diverse organisms to infer ecological and evolutionary outcomes. These TPCs are typically characterized by estimating a trait (e.g. bite rate, mortality rate, development rate) across a gradient of constant temperatures in a controlled laboratory study^13^. For modeling mosquito-borne diseases, the TPCs are then often incorporated either into standard formulae for the pathogen’s basic reproduction number (*R_0_*; defined as the number of secondary cases arising from a primary case introduced into a fully susceptible population) to predict overall thermal suitability for transmission^2,8,9,19^, or into mechanistic dynamical models used to predict human incidence or the final epidemic size^20,21^. These approaches have generated many important insights, including: 1) warming at northern latitudes or high elevations has increased and will continue to increase suitability for transmission due to longer and more intense transmission seasons, resulting in the potential for large epidemics^3,7^; 2) areas of the world that are currently suitable for transmission may become less environmentally suitable as temperatures warm beyond the optimum^3,22^; 3) disease intervention efforts (e.g. vector control, vaccination, drug coverage) will need to be more expansive in areas of the world and times of year that are most suitable for transmission^3,23,24^; and 4) climate change could shift disease burden from malaria to arboviruses in Africa ^25^.

Although controlled laboratory studies have provided insight into the effects of temperature on mosquito life history, mosquito-pathogen interactions, and overall transmission potential, the temperatures that organisms experience in the field are highly variable, fluctuating diurnally, seasonally, annually, and on other timescales ^26^. A consequence of the mathematical fact known as ‘Jensen’s inequality’ is that when temperature impacts performance non-linearly, then the time-average of performance across a thermally fluctuating environment is not equal to the performance measured at the average temperature^27–29^. Specifically, thermal fluctuations should increase performance in accelerating (convex) portions of a TPC and decrease performance in decelerating (concave) portions (**Figure 1B**), simply based on the time spent at each temperature and the associated performance predicted by the TPC. This theoretical prediction is supported by a growing body of empirical research across diverse taxa^30–34^, including mosquitoes^35–37^, demonstrating that trait performance under fluctuating temperatures can differ substantially from performance at constant temperatures.

**Figure 1:**
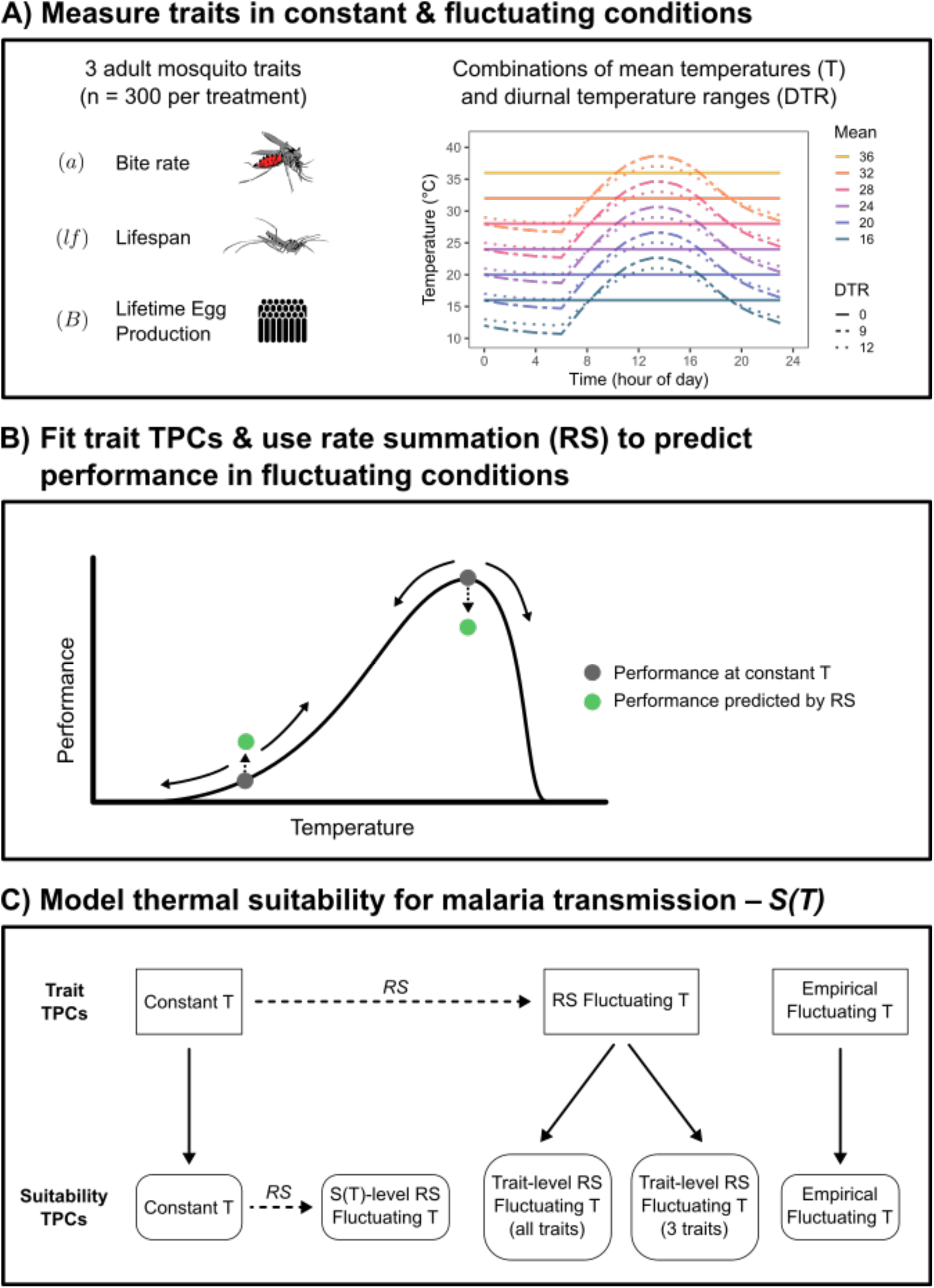
Conceptual figure summarizing the study. A) We measured three adult mosquito traits (bite rate [a], lifespan [lf], and lifetime egg production [B]) in constant and fluctuating conditions (diurnal temperature range [DTR] = 0, 9, and 12°C) across a range of mean temperatures (mean temperatures = 16, 20, 24, 28, and 32°C for all DTR treatments; 36°C for DTR = 0°C only). B) For each trait, we fit thermal response curves (TPCs) to the data from each DTR treatment. Additionally, we used rate summation (RS) to predict performance in fluctuating environments based on the TPC fitted to data from constant environments. Compared to constant temperatures with the same mean (dark gray points), in fluctuating temperatures (solid arrows) rate summation will predict a decrease in performance over decelerating portions of a TPC (e.g., near the optimum) and an increase in performance over accelerating portions of a TPC (dashed arrow and green points). C) We compared five versions of a model predicting thermal suitability for transmission, S(T), parameterized with different trait TPCs. Model 1 (‘Constant T’) used TPCs fit to trait data from constant temperatures. Model 2 (‘Empirical Fluctuating T’) used TPCs fit to trait data from fluctuating temperatures. Models 3 and 4 used TPCs generated by applying rate summation to constant temperature TPCs for either the 3 focal traits measured here (model 3: ‘Trait-level RS Fluctuating T - 3 traits’) or all traits in the model (model 4: ‘Trait-level RS Fluctuating T - all traits’). Model 5 (‘S(T)-level RS Fluctuating T’) applied rate summation directly to the TPC for suitability generated in model 1. Dashed arrows denote RS calculations and solid arrows denote parameterizing the suitability model with trait TPCs.

Unfortunately, it is logistically infeasible to experimentally evaluate every possible fluctuating temperature regime that an organism might experience. Accordingly, studies typically use ‘non-linear averaging’ or ‘rate summation’ to quantitatively predict the average trait performance as temperature fluctuates over time^27,30,38–41^ (**Equation 1**).

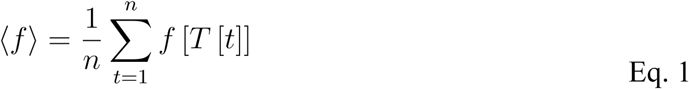

Here,

<*f*> is the average performance of a trait, and is calculated from *f*, the trait performance as a function of temperature (*T*), which in turn is a function of time (*t*) from *t=*1 to *t=n*. This approach has been adopted widely to account for the impact of temperature variation on the thermal suitability of transmission in many vector-borne disease systems^10,35,42–48^.

However, rate summation makes two simplifying assumptions that are likely to be violated in many biological systems: 1) traits always exhibits the same value at a given temperature in both fluctuating and constant environments; and 2) performance changes instantaneously with temperature (i.e., no acclimation period). First, performance in fluctuating environments can differ from in the equivalent constant temperatures due to the inherent effects of thermal fluctuations on organismal performance^49^, including acclimation to thermal stress^13,50,51^, accumulation of damage associated with thermal stress^52,53^, and processes to repair damage incurred at extreme low or high temperatures during time spent at more favorable temperatures (e.g. production of heat shock proteins) that cause hysteresis effects^51,54,55^. Second, time lags and other thermal acclimation effects are common and varied in their impact on performance^56^. Although the accuracy of rate summation has been assessed in other organisms^28,30,39,49^, it has not been evaluated for vector-borne disease systems. Furthermore, because rate summation has mixed success in predicting performance of traits, it is unclear whether rate summation can accurately predict suitability for mosquito-borne disease transmission. Evaluating the ability of rate summation to capture the thermal suitability of realistically fluctuating conditions has important implications for understanding how mosquito populations and their transmission dynamics will play out in natural field settings, as well as in response to future climate change.

In this study, we use experimental data and modeling (**Figure 1**) to better understand the use of rate summation to predict the thermal suitability for malaria transmission by *Anopheles stephensi,* an important mosquito vector of urban malaria in South Asia and now Africa. Specifically, we ask: 1) Do field-relevant diurnal temperature fluctuations alter the relationships between temperature and adult mosquito life history traits compared to those characterized across constant temperatures? 2) Can rate summation accurately predict these temperature-trait relationships in environments that diurnally fluctuate? 3) How do these various temperature-trait relationships scale up to impact predicted thermal suitability for malaria transmission? Our results show that temperature fluctuations significantly alter the thermal responses of adult mosquito traits, that rate summation largely fails to predict the performance of these traits, and that this discrepancy impacts the predicted thermal limits for malaria transmission. We discuss reasons for why rate summation might fail to predict performance in a fluctuating environment and the implications for using this technique in mechanistic modeling frameworks that predict vector-borne disease transmission.

## Study and Suitability Model Overview

We modeled the effects of diurnal temperature fluctuation on predicted thermal suitability for transmission of malaria, *S(T)*, using a trait-based mechanistic model based on a standard derivation of *R_0_* for malaria (**Equation 2**, see *Methods*). We focused on the impacts of three adult mosquito traits that we directly measured across temperature gradients in both constant and fluctuating conditions: daily female bite rate (*a*), lifetime egg production (*B*), and lifespan (*lf*). Data for other traits required to calculate *S(T)*–larval survival (*p_EA_*), development rate (*MDR*), vector competence (*bc*), and extrinsic incubation period (*EIP*)–were taken from previous experimental studies with constant temperature gradients^19,36^. Trait thermal performance curves (TPCs) were fitted using either a symmetric (quadratic) or asymmetric (Brière) function, chosen by comparing Deviance Information Criterions (DIC)^67^.

We generated five versions of the *S(T)* model (**Figure 1**) parameterized with TPCs for traits either fit to data from three different temperature fluctuation regimes (diurnal temperature range [DTR] = 0, 9, or 12°C) or calculated via rate summation (RS).

1. TPCs fit to trait data from across a range of constant temperatures (‘constant’).
2. TPCs fit to trait data from fluctuating conditions for the focal traits with empirical data (*a, B,* and *lf*), combined with TPCs fit to trait data from constant temperatures for traits measured in other studies (*p_EA_, MDR, bc,* and *EIP*; ‘empirical fluctuating’).
3. TPCs generated by applying rate summation to the TPCs from constant temperatures for the focal traits (*a, B,* and *lf*); as in version 2, other traits (*p_EA_, MDR, bc,* and *EIP*) used unmodified TPCs from constant temperatures (‘trait-level RS - 3 traits’).
4. Similar to version 3 above, but rate summation was applied to the TPCs from constant temperatures for all traits (‘trait-level RS - all traits’).
5. Rate summation applied to the TPC for *S(T)* generated from traits measured across a range of constant temperatures (i.e, the output of constant model 1 above) to generate a new TPC for *S(T)* (‘*S(T)*-level RS’).

We used these five versions of *S(T)*, generated for both fluctuating DTRs (9 and 12°C) where applicable, to assess the following questions: A) how thermal suitability is likely affected by temperature fluctuations (model 1 versus model 2); B) if rate summation can adequately predict suitability in fluctuating temperature regimes (model 2 versus model 3); and C) how the level at which rate summation is calculated (on the component traits or on suitability itself) impacts predictions (model 4 versus model 5).

## Results

### Diurnal fluctuation alters the thermal responses of mosquito traits

All three focal traits (bite rate [*a*], lifespan [*lf*], and lifetime egg production [*B*]) responded strongly to mean temperature (**Figure 2**). The shape of the thermal response was relatively consistent for each trait across fluctuation treatments (diurnal temperature range [DTR] = 0, 9, or 12°C). Lifespan (*lf*) always responded symmetrically and was best fit with a quadratic function, while bite rate (*a*) always responded asymmetrically and was best fit with a Brière function. Lifetime egg production (*B*) was fit similarly by both functions (ΔDIC < 2.0 for all fluctuation treatments); we elected to always use a quadratic function to be consistent and because it had a slightly lower DIC for two of three fluctuation treatments (**Table S1**).

**Figure 2:**
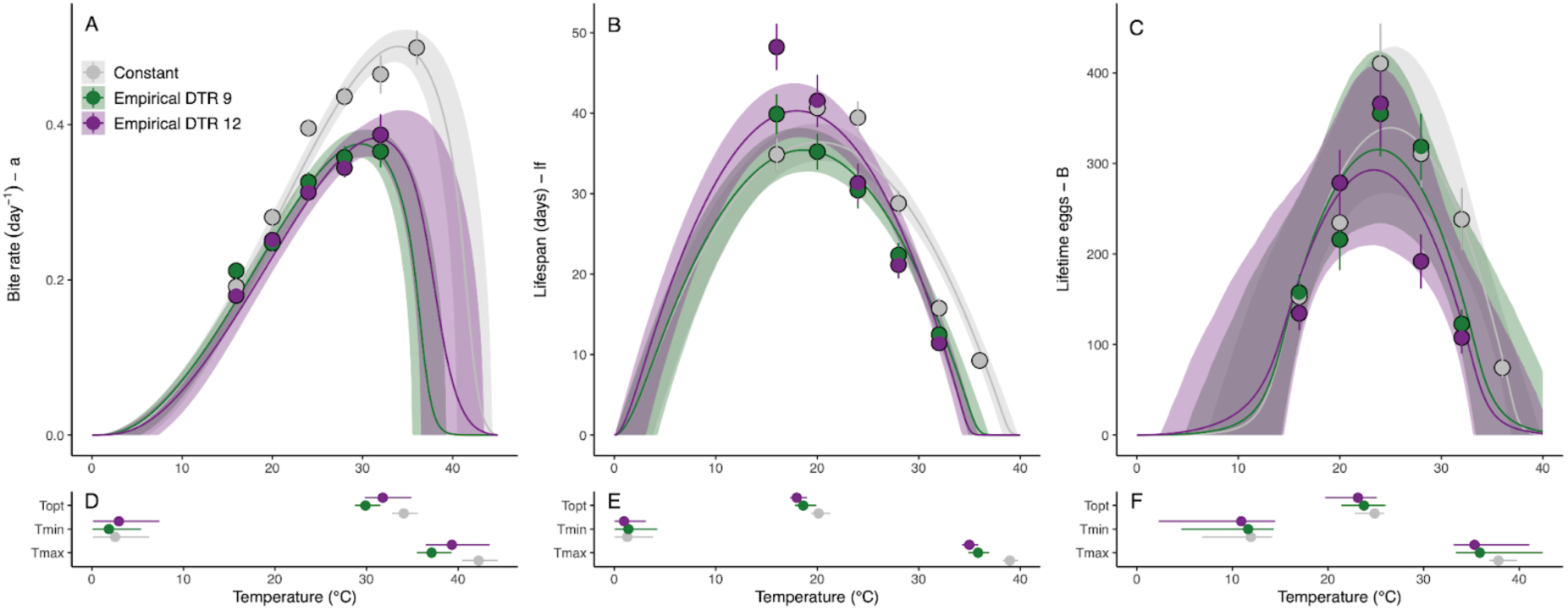
Empirically measured thermal responses for three adult *Anopheles stephensi* traits in constant and diurnally fluctuating temperatures. Traits include: bite rate (a, left column), lifespan (lf, center column) and lifetime egg production (B, right column). Colors denote daily temperature range (DTR) treatment: 0°C (gray), 9°C (green), and 12°C (purple). A-C) Summarized data and thermal performance curves (TPCs). TPC contours show posterior distribution medians, with 95% credible intervals as shaded areas. Points and error bars display block means and standard errors for visual comparison between treatments. (TPCs were fit to individual-level data.) D-F) Key temperature values from the TPCs: thermal optimum (Topt), thermal minimum (Tmin), and thermal maximum (Tmax). Points display posterior distribution medians and error bars display 95% credible intervals.

Fluctuating temperatures significantly altered the thermal performance curves (TPCs) for each trait (**Figure 2, Tables 1, S2, and S3**). These changes were reflected by shifts in TPC characteristics as well as the magnitude of performance in each environment. Diurnal temperature fluctuations caused downward shifts in three key TPC parameters (*T_opt_*, *T_max_*, and *T_breadth_*) and the magnitude of these shifts depended on the trait. For all three parameters, the shifts were largest for bite rate (*a*), followed by lifespan (*lf*), and then lifetime egg production (*B*). TPCs characterized under temperature fluctuations resulted in cooler predicted thermal optima (*T_opt_*), ranging from 1.2-4.2°C cooler, and thermal maxima (*T_max_*), ranging from 2.5-5.2°C cooler, depending on the trait. We were unable to detect any shifts in the thermal minima (*T_min_*). Consequently, we also observed a narrowing in thermal breadth (*T_breadth_*) that ranged from from 2.3-4.5°C depending on the trait. Differences in TPCs based on the magnitude of fluctuation (i.e., DTR 9°C vs. DTR 12°C) were only significant for lifespan (*lf*).

**Table 1.**
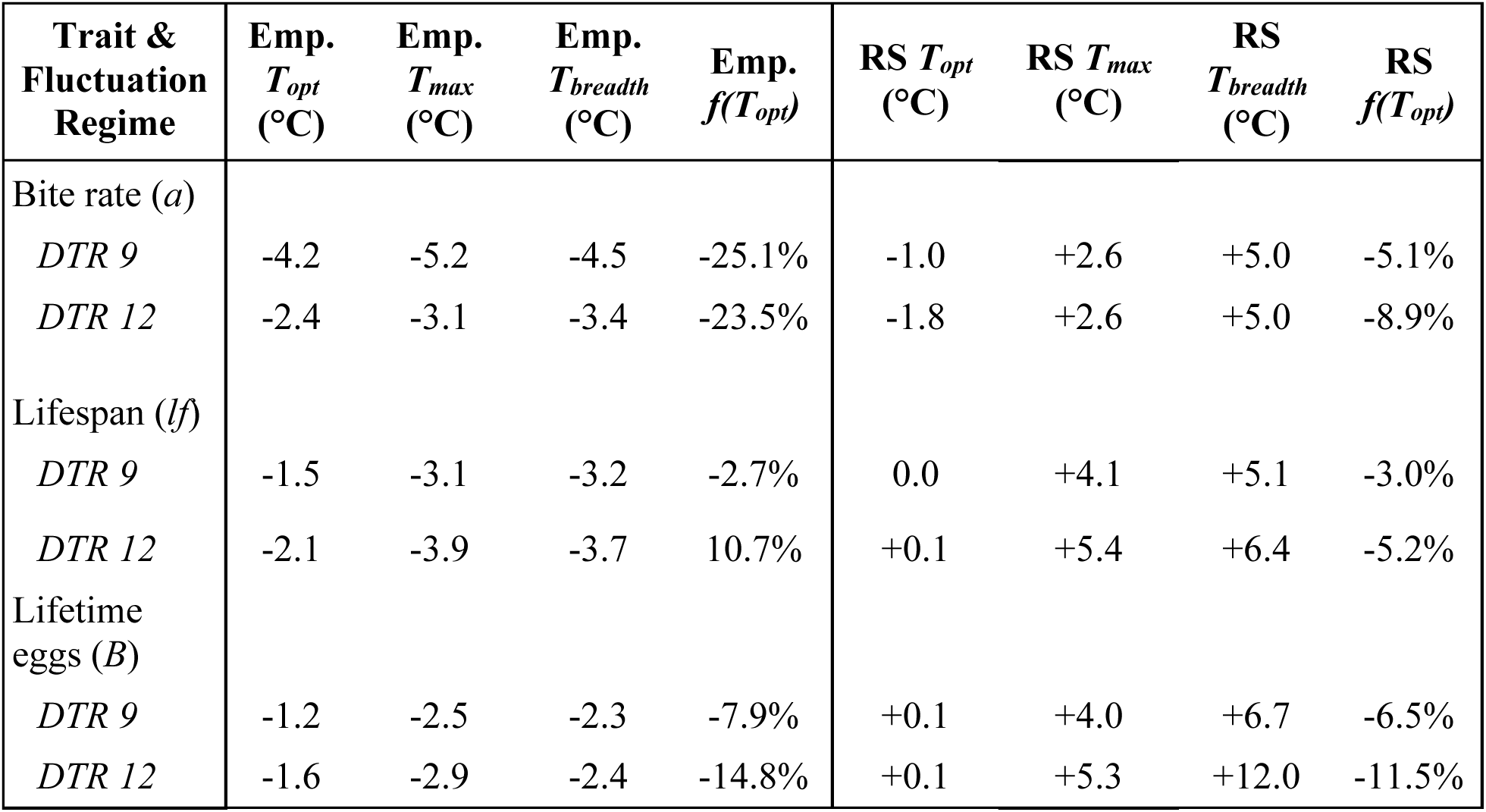
Shifts in properties of thermal performance curves (TPCs) for adult mosquito traits due to temperature fluctuations. Differences in thermal optimum (*T_opt_*), thermal maximum (*T_max_*), and thermal breadth (*T_breadth_*), and percent change in the predicted trait value at *T_opt_* [*f(T_opt_)*]. TPCs fit to empirical data from fluctuating temperatures (Emp.) and TPCs calculated using rate summation (RS) are both compared to TPCs fit to data from constant temperatures. Diurnal temperature ranges (DTR) = 9 and 12°C. Differences calculated using median values. See **Table S2** for the original parameter values for each model.

Fluctuating temperatures also decreased absolute performance for all traits at their thermal optima and warmer temperatures, relative to trait performance at constant temperatures (**Figure 2**, **Table 1**). Maximum predicted performance [i.e., trait value at the thermal optimum, *f(T_opt_)*] decreased more for bite rate (*a*; 23.5-25.1% lower) than for lifespan (*lf*; 2.7% lower to 10.7% higher) or egg production (*B*; 7.9-14.8% lower). For lifespan (*lf*), fluctuating temperatures increased performance relative to constant temperatures at 16°C, which increased the maximum predicted performance for DTR 12°C only.

### Rate summation fails to predict thermal responses in fluctuating environments

Overall, rate summation failed to accurately predict trait performance in a diurnally fluctuating thermal environment. Rate summation did not predict the observed shifts in three TPC parameters (*T_opt_*, *T_max_*, and *T_breadth_*) or the maximum predicted performance (**Figure 3**, **Table 1**). In fact, for two of the parameters (*T_max_*, and thus also *T_breadth_*), rate summation predicted that temperature fluctuations would change performance in a different direction than what was observed (i.e., it predicted warmer/wider shifts instead of cooler/narrower shifts relative to performance in constant temperature conditions). Rate summation also predicted small decreases in *T_min_* under fluctuations, which we did not detect in the TPCs fit to empirical data from fluctuating conditions.

**Figure 3:**
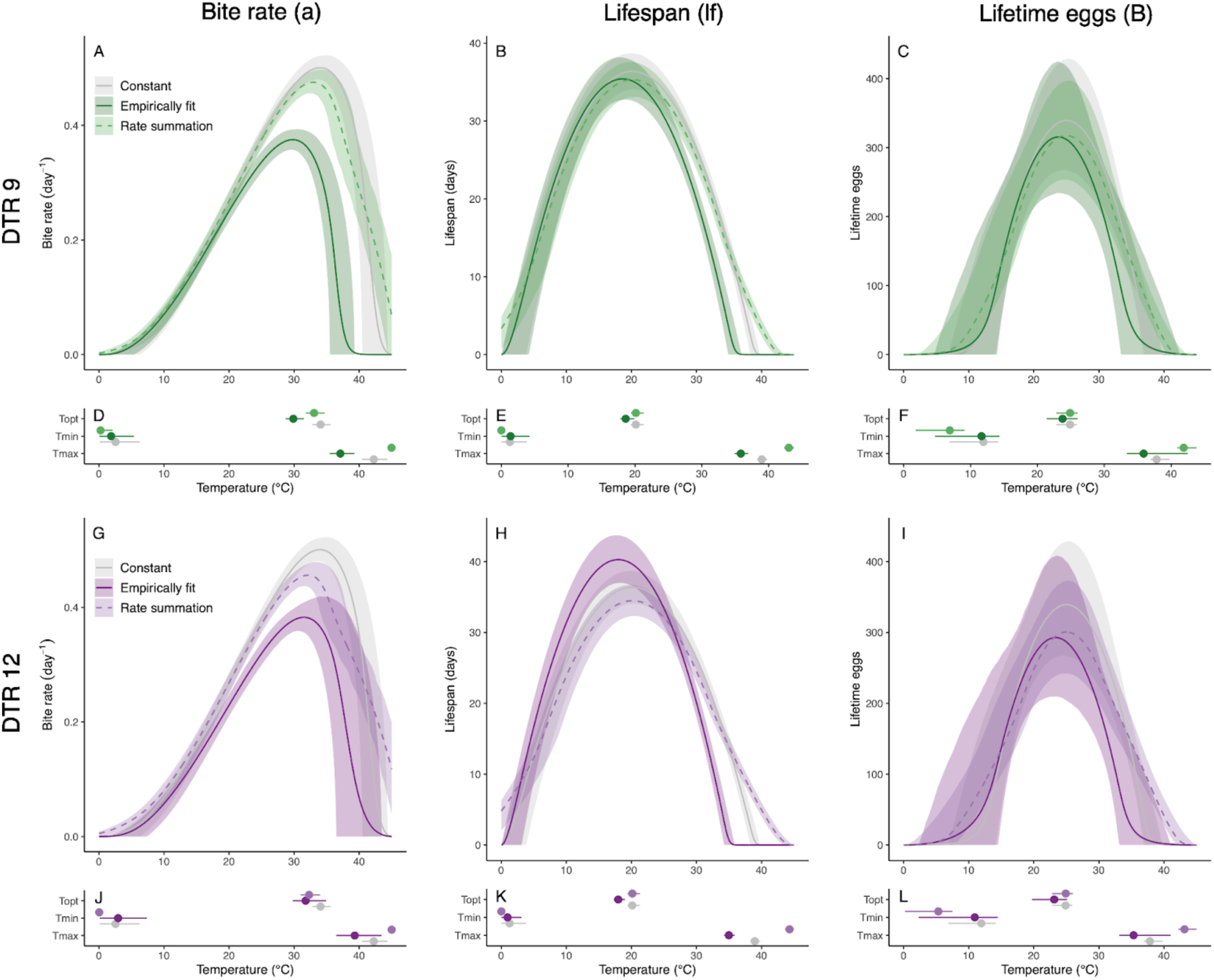
Thermal performance based on empirical observations or predictions generated by rate summation for *Anopheles stephensi* performance in diurnally fluctuating temperature environments. Left column: bite rate (a), center column: lifespan (lf), right column: lifetime egg production (B). Top row (green): daily temperature range (DTR) 9°C, bottom row (purple): DTR 12°C. Darker hues and solid lines show thermal performance curves (TPCs) fit to empirical data collected from mosquitoes housed in diurnally fluctuating temperature conditions. Light hues and dashed lines show predictions generated by rate summation. TPCs fit to empirical data collected from mosquitoes housed across constant temperature conditions shown in gray for comparison.

Rate summation overestimated the *T_max_* for bite rate (*a*), lifespan (*lf*), and to some degree lifetime egg production (*B*, for DTR 12°C) (**Figure 3**, **Table 1**). It predicted increases in the thermal maxima (*T_max_*) for all three traits relative to mosquitoes housed in constant temperatures (2.6-5.4°C warmer). In contrast, mosquitoes housed in fluctuating conditions had cooler *T_max_* for all traits relative to those housed under constant temperature conditions (2.5-5.2°C cooler). As a result, rate summation overpredicted the overall thermal breadth (*T_breadth_*) of trait performance relative to mosquitoes housed in constant temperatures (5.0-12.0°C warmer), instead of the more constrained thermal breadth observed for mosquitoes in thermally fluctuating environments (2.3-4.5°C cooler).

Rate summation generally overestimated the *T_opt_* in mosquitoes housed in fluctuating environments and failed to predict differences compared to those housed in constant temperature conditions (**Figure 3**, **Table 1**). This trend was strongest (minimal overlap in credible intervals) for the daily bite rate (*a*) at DTR 9°C and lifespan (*lf*) for both DTR 9°C and 12°C treatments. For bite rate (*a*), rate summation underestimated the decrease in the *T_opt_* that was observed in mosquitoes housed under fluctuating thermal conditions relative to constant temperature conditions (1.0-1.8°C cooler predicted by rate summation vs. 2.4-4.2°C cooler from empirical data). For lifespan (*lf*) and lifetime egg production (*B*), rate summation predicted essentially no change in the *T_opt_* from mosquitoes housed at constant temperatures, in contrast to observed decreases in the *T_opt_* in mosquitoes housed under temperature fluctuations (1.2-2.1°C cooler).

In many cases, rate summation also failed to accurately predict absolute trait performance in fluctuating environments (**Figure 3**, **Table 1**). In the most extreme example, for daily bite rate (*a*), rate summation predicted substantially higher maximum trait performance [*f(T_opt_)*] for both DTR treatments (predictions 16.1-21.1% higher than empirical observations). For lifetime egg production (*B*) and lifespan (*lf*) in DTR 9°C, rate summation was fairly accurate at predicting small decreases in maximum trait performance [*f(T_opt_)*; predictions all within 3.7% of empirical observations]. However, for lifespan (*lf*) in DTR 12°C, rate summation predicted small decreases in absolute trait performance at cooler temperatures, when TPCs fit to observations yielded increases in absolute trait performance relative to constant temperatures.

### Diurnal temperature fluctuation impacts the predicted suitability for transmission

The effects of fluctuating temperatures on the three adult mosquito traits measured here lowered the predicted suitability for transmission, *S(T)*, at warmer temperatures (**Figure 4A**, **Table 2**). As a result, model 2 (empirical fluctuating) lowered the predicted *T_opt_* by 1.2-1.4°C, *T_max_* by 0.8-1.8°C, thermal breadth by 0.8-1.2°C, and predicted amount of suitability at *T_opt_* by 32.0-33.8% (**Table 2**) compared to model 1 (constant). Applying rate summation to the trait TPCs to predict performance of the three adult mosquito traits in thermally fluctuating environments (model 3: trait-level RS - 3 traits) did not capture these effects (**Figure 4B**). This model predicted much smaller changes in *T_opt_* (0.1-0.2°C lower), no change in *T_max_* or the thermal breadth, and smaller reductions in suitability at *T_opt_* (10.0-17.1% lower) (**Table 2**), therefore overestimating suitability near and above the thermal optimum compared to model 2 based on empirical observations. Finally, the level at which the rate summation calculation was performed (on all seven traits prior to calculating suitability [model 4: Trait-level RS - all traits] or directly on the suitability curve [model 5: *S(T)*-level RS]) visually impacted the curves for predicted suitability (**Figure 4C**), but had little impacts on the key values of the TPCs (*T_min_, T_opt_, T_max_, T_breadth_*) or the predicted reduction in suitability at *T_opt_* (18.1-32.0% lower) (**Table 2**). Performing rate summation on the *S*(*T*) curve yielded a TPC that was wider and predicted higher suitability at temperatures near the thermal margins (**Figure 4C**). However, this difference was not reflected in the values for *T_min_* or *T_max_*, which were identical for a given level of diurnal temperature variation (DTR), because the suitability curves for model 5 approached the x-axis extremely gradually. Additionally, the predicted optimum (*T_opt_*) and the magnitude of transmission near the optimum was very similar for both versions of suitability (Trait-level RS: 0.2-0.4°C cooler than constant temperatures, *S*(*T*)-level RS: 0.1°C warmer or cooler than constant temperatures; **Figure 4C**, **Table 2**).

**Figure 4:**
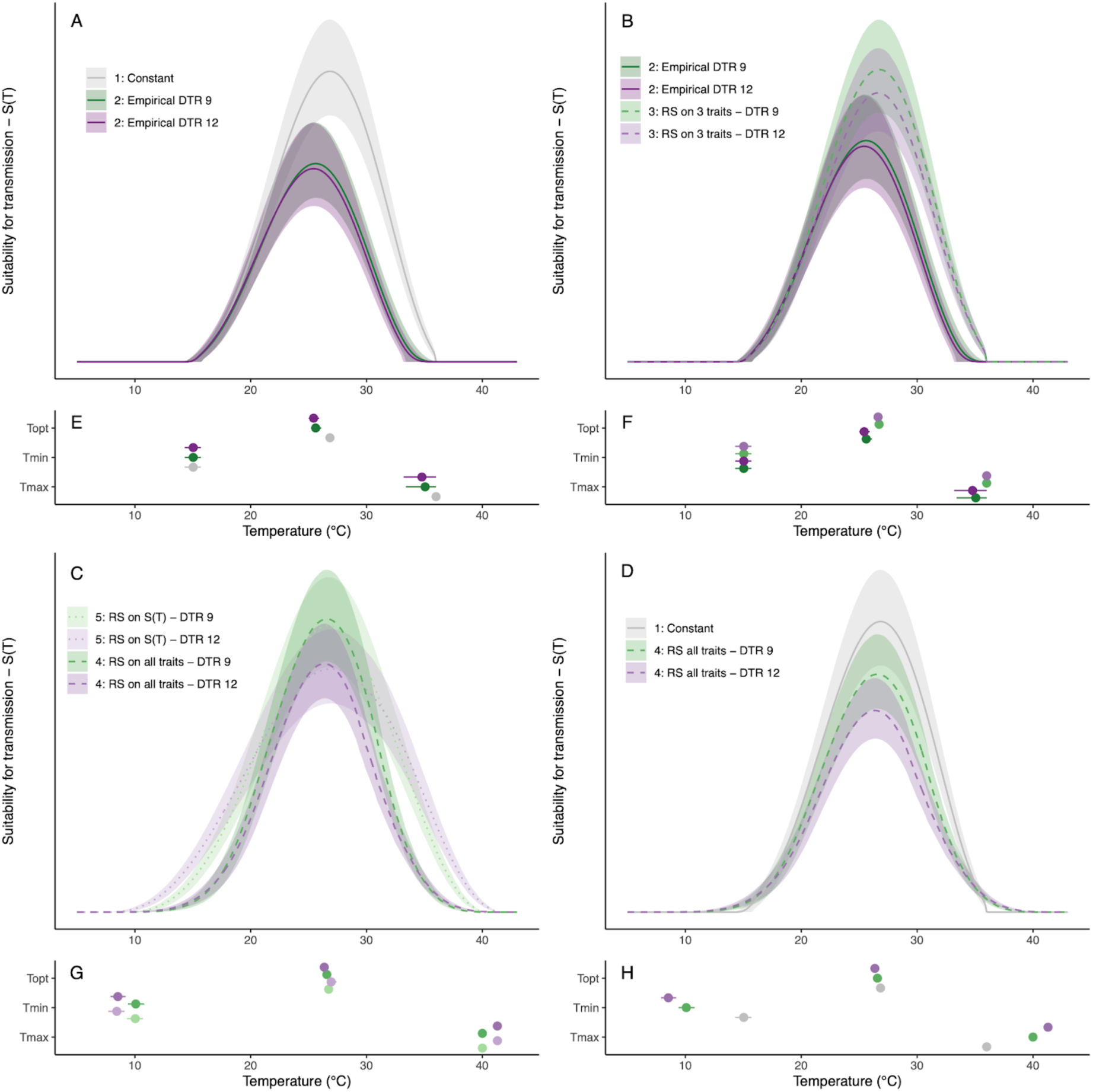
Thermal suitability for transmission of malaria by *Anopheles stephensi* predicted for constant and diurnally fluctuating temperature conditions. A) Model versions parameterized with TPCs fit to empirical data collected from constant temperature (T) conditions (model 1, constant T) and fluctuating conditions (model 2, empirical fluctuating T). B) Model versions parameterized with TPCs fit to empirical data collected from fluctuating conditions (model 2, empirical fluctuating T) and TPCs predicted by rate summation performed on trait TPCs for focal traits only (model 3, trait-level RS - 3 traits). C) Model versions comparing rate summation performed on the TPCs for traits (model 4, trait-level RS - all traits) and on the TPC for suitability itself (model 5, *S(T)*-level RS). The numbers in the legends below refer to model numbers, see *Methods* for model details.

**Table 2:** Shifts in properties of thermal performance curves (TPCs) for models of predicted suitability of malaria transmission, *S*(*T*), due to temperature fluctuations. Differences in thermal optimum (*T_opt_*), thermal maximum (*T_max_*), and thermal breadth (*T_breadth_*), and the percent change in median *S*(*T*) predicted at *T_opt_*, compared to the constant temperature model (model 1). Fluctuating models are parameterized with trait TPCs fit from empirical data (model 2: “Empirical fluctuating”) or are calculated using rate summation (RS). Rate summation was used only for the three traits with empirical data (model 3: “Trait-level RS - 3 traits”), for all traits (model 4: “Trait-level RS - all traits”), or directly on the TPC for suitability, *S*(*T*), at constant temperatures (model 5: “*S*(*T*)-level RS”). Diurnal temperature ranges (DTR) = 9 and 12°C. Differences calculated using median values. See **Table S3** for original parameter values for each model.

| Model & Fluctuation Regime | $T_{opt}$ (°C) | $T_{max}$ (°C) | $T_{breadth}$ (°C) | $S(T_{opt})$ |
| --- | --- | --- | --- | --- |
| 2) Empirical fluctuating - 3 traits |  |  |  |  |
| *DTR 9* | -1.2 | -0.8 | -0.8 | -32.0% |
| *DTR 12* | -1.4 | -1.8 | -1.2 | -33.8% |
| 3) Trait-level RS - 3 traits |  |  |  |  |
| *DTR 9* | -0.1 | 0.0 | 0.0 | -10.0% |
| *DTR 12* | -0.2 | 0.0 | 0.0 | -17.1% |
| 4) Trait-level RS - all traits |  |  |  |  |
| *DTR 9* | -0.2 | +4.0 | +9.0 | -18.1% |
| *DTR 12* | -0.4 | +5.3 | +11.9 | -30.6% |
| 5) $S(T)$-level RS | | | | |
| *DTR 9* | -0.1 | +4.0 | +9.0 | -19.9% |
| *DTR 12* | +0.1 | +5.3 | +11.9 | -32.0% |

The sensitivity and uncertainty analyses provide insight into which traits determine key characteristics of the TPC for suitability (*T_min_*, *T_opt_*, and *T_max_*) and drive uncertainty across the temperature gradient (**Figures S1, S2** and **S3**). For all suitability models, as temperature increases, lifespan (*lf*) is most important for lowering *T_opt_* while bite rate (*a*) and development rate (*MDR*) are most important for raising *T_opt_* (**Figures S1** and **S2**). Together, these traits most strongly influence the optimal temperature for transmission (*T_opt_*), consistent with previous studies^4^. In model 1 (constant), *T_min_* and *T_max_* are both determined by larval traits not measured in this study (larval survival [*p_EA_*] and development rate [*MDR*], respectively). The TPC for development rate [*MDR*] has very little uncertainty in its *T_max_*, which leads to similarly low uncertainty for the *T_max_* of suitability. Most of the uncertainty in model 1 is generated by lifetime egg production (*B*) near *T_opt_* and by vector competence (*bc*) near both thermal margins (**Figure S3**).

By contrast, in model 2 (empirical fluctuating), *T_max_* for suitability is determined primarily by the effects of temperature on mosquito lifespan (*lf*), and then lifetime egg production (*B*), as the *T_max_* for both of those traits decrease below the *T_max_* for development rate (MDR; **Figures S1** and **S2**). Larval survival (*p_EA_*) still determines *T_min_* and uncertainty in vector competence and lifetime egg production (*B*) are still most important near the lower thermal limit and optimum, respectively. However, near the upper thermal limit, most of the uncertainty is now due to lifetime egg production (*B*) and lifespan (*lf*; **Figure S3**). Model 3 (trait-based RS - 3 traits) retains the effects of the unmodified TPCs for larval survival (*p_EA_*) and development rate (*MDR*) from model 1 (constant), which again determine *T_min_* and *T_max_*, respectively (**Figures S1** and **S2**). Models 4 (trait-level RS - all traits) and 5 (S(T)-level RS) preserve the importance of these two larval traits for determining *T_min_* and *T_max_*, but the rate summation calculation changes the specific temperature at which *T_min_* and *T_max_* occur. Models 3, 4, and 5 also retain the uncertainty patterns from model 1: lifetime egg production (*B*) is most important near *T_opt_* and vector competence (*bc*) is most important near both thermal margins (**Figure S3**).

### Mapping predicted suitability for transmission

Differences in the predicted thermal suitability can be visualized on maps showing the number of months predicted to have temperatures suitable for transmission, *S*(*T*) > 0.001, in both the native zone (Central and South Asia; **Figure 5** left column) and introduced zone (Africa; **Figure 6** left column) for *An. stephensi*. The constant temperature model for suitability (model 1), predicts that India is suitable for malaria transmission year round (**Figures 5A**), as is much of Africa **(Figure 6A**). The empirical fluctuations model for suitability (model 2) shows a slightly shorter transmission season in Northern India and Pakistan (**Figure 5B**) and in Northern Africa (**Figure 6B**), due to its cooler *T_max_* value (**Table 3**). Both suitability models based on rate summation calculations (model 4: trait-level RS and model 5: *S*(*T*)-level RS) yielded *T_min_* values that were much cooler than models 1 and 2 (**Table 3**), and thus produced maps with predicted year-round transmission across all of India (**Figure 5C-D)** and nearly all of Africa (**Figure 6C-D**). Compared to the constant and empirical fluctuating models, both rate summation models predicted much longer transmission seasons in Northern India, Pakistan, and Iran (**Figure 6C-D**), as well as in Northern and Southern Africa (**Figure 7C-D**). Overall, the predictions from the constant temperature model were more like those from the empirical fluctuating model, while the predictions from both rate summation models diverged more (**Figures 5 and 6** left columns, **Table 3**).

**Figure 5:**
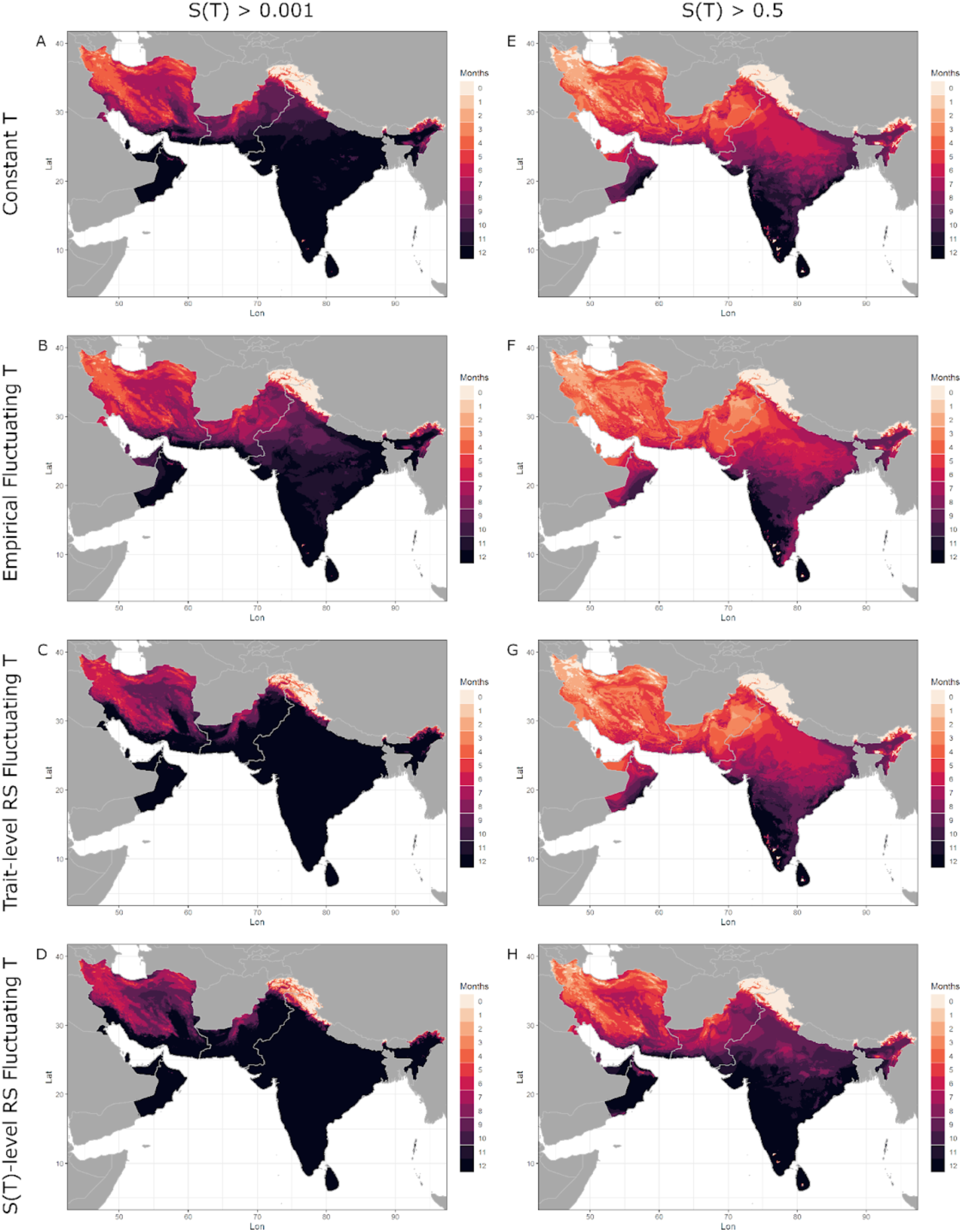
Months of thermal suitability, S(T), for transmission of malaria by *Anopheles stephensi* in its native range in Central and South Asia predicted by models parameterized using constant and fluctuating temperatures. Left column: total months where *S(T)* is predicted to exceed 0.001 (i.e. when transmission is possible). Right column: total months where *S(T)* is predicted to exceed 0.5 (i.e. when transmission is relatively favored by temperature). Darker hues indicate more months. Top row: model 1 (constant T) uses trait TPCs fit to data across a range of constant temperatures; second row: model 2 (empirical fluctuating T) uses trait TPCs fit to data across a range of fluctuating temperatures; third row: model 4 (trait-level RS - all traits), uses trait TPCs generated by applying rate summation to TPCs fit to data from constant temperatures for all traits; bottom row: model 5 (*S(T)*-level RS), applies rate summation to the TPC for suitability generated from traits measured across a range of constant temperatures (i.e. the output of version 1). Fluctuating temperature models used a daily temperature range (DTR) = 12°C.

**Figure 6:**
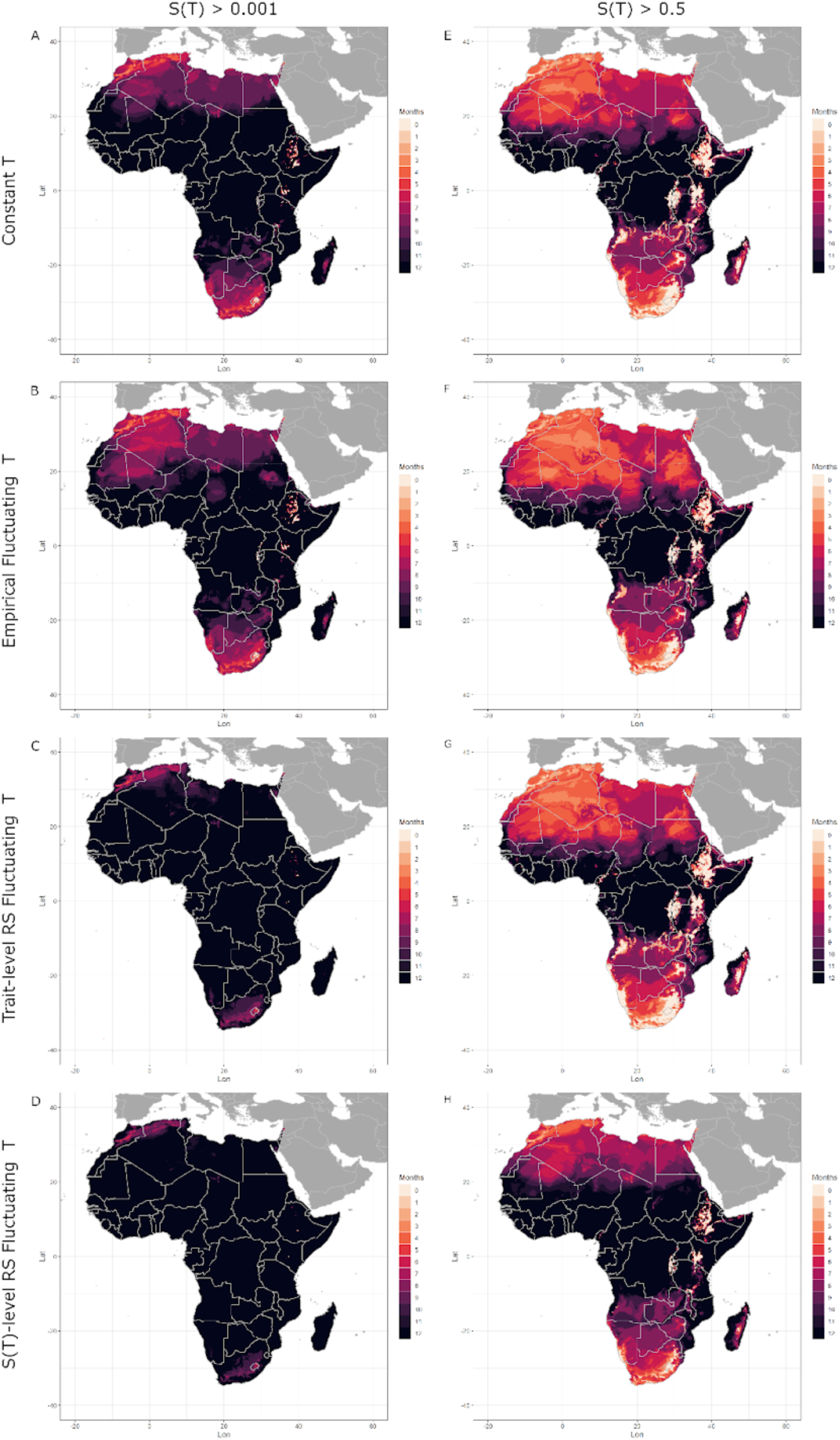
Months of thermal suitability, S(T), for transmission of malaria by *Anopheles stephensi* in its invading range in Africa predicted by models parameterized using constant and fluctuating temperatures. Left column: total months where *S(T)* is predicted to exceed 0.001 (i.e. when transmission is possible). Right column: total months where *S(T)* is predicted to exceed 0.5 (i.e. when transmission is relatively favored by temperature). Darker hues indicate more months. Top row: model 1 (constant T) uses trait TPCs fit to data across a range of constant temperatures; second row: model 2 (empirical fluctuating T) uses trait TPCs fit to data across a range of fluctuating temperatures; third row: model 4 (trait-level RS - all traits), uses trait TPCs generated by applying rate summation to TPCs fit to data from constant temperatures for all traits; bottom row: model 5 (*S(T)*-level RS), applies rate summation to the TPC for suitability generated from traits measured across a range of constant temperatures (i.e. the output of version 1). Fluctuating temperature models used a daily temperature range (DTR) = 12°C.

**Table 3:** Temperature thresholds used for mapping four models of thermal suitability. Four versions of the model for thermal suitability, *S*(*T*), parameterized with different trait TPCs or calculated using rate summation (RS): trait TPCs fit from empirical data under constant temperatures (model 1: “Constant”), trait TPCs fit from empirical data under fluctuating temperatures (model 2: “Empirical Fluctuating”), RS at the trait-level for all traits (model 4: “Trait-level RS - all traits”), or RS directly on the TPC for suitability, *S*(*T*), parameterised under constant temperatures (model 5: “*S*(*T*)-level RS”). All fluctuating models were for Diurnal temperature ranges (DTR) = 12°C only. Units are in °C.

| Model | Range where $S(T) > 0.001$ (°C) | Range where $S(T) > 0.5$ (°C) |
| --- | --- | --- |
| Constant (1) | 15.8 - 35.8 | 21.1 - 31.9 |
| Empirical Fluctuating (2) | 15.8 - 33.1 | 20.3 - 30.2 |
| Trait-level RS - all traits (4) | 10.9 - 40.1 | 21.2 - 31.1 |
| $S(T)$-level RS (5) | 9.3 - 41.1 | 18.7 - 34.7 |

Both suitability models based on rate summation calculations (model 4: Trait-level RS and model 5: *S*(*T*)-level RS) yielded nearly identical results for *S*(*T*) > 0.001 (**Figures 5 and 6**). However, even though the *T_min_* and *T_max_* of thermal suitability is predicted to be the same across both models, there are clearly differences in the rate at which temperatures increase from or decrease toward the *T_min_* and *T_max_*, respectively, across models. When we use a higher threshold *S*(*T*) > 0.5, for where the thermal suitability is relatively high (**Figures 5 and 6** right columns), performing rate summation on the TPC for suitability (model 5, **Figure 5D and 6D**) predicts more areas with relatively high thermal suitability year-round than performing rate summation on the TPCs of the component traits (model 4, **Figure 5C and 6C**). For this higher threshold, performing rate summation at the trait-level produced maps that were quite like both empirical models (constant and fluctuating), while performing rate summation directly on the suitability TPC did not (**Figures 5 and 6** right columns, **Table 3**).

## Discussion

This study measured and analyzed adult mosquito life history traits (lifespan, bite rate, and lifetime reproductive output) for the urban Asian malaria vector *Anopheles stephensi* across a temperature gradient under three daily temperature range (DTR) regimes (0, 9, and 12°C). We used these data to determine if standard modeling techniques could accurately predict the impact of biologically relevant daily temperature fluctuations on mosquito performance and environmental suitability for malaria transmission. We found that: 1) daily temperature fluctuation significantly altered the thermal responses for these critical mosquito traits involved in pathogen transmission; 2) rate summation (RS), a non-linear averaging approach used to estimate the effect of temperature fluctuations using thermal performance curves (TPCs) characterized in constant temperature environments, did not accurately predict trait thermal responses in diurnally fluctuating temperature environments; and 3) while thermal suitability predictions constructed with responses from constant temperature conditions did not capture the impact of real-world temperature variation on mosquito traits, they were substantially more accurate for predicting and mapping the thermal limits of malaria transmission than predictions constructed using rate summation calculations. This result stems from a general property of performing rate summation on TPCs that cut-off at the x-axis, as is often the case for biological traits that cannot take negative values. Thus, we conclude that while daily-scale temperature fluctuations have important impacts on organismal performance, for some applications it may be better to use thermal responses fit under constant temperature environments than to try to incorporate the impact of fluctuating temperatures using non-linear averaging. Additionally, it is vital to improve methods of estimating the physiological effects of temperature fluctuation in real-world situations to accurately predict the thermal suitability for transmission of vector-borne diseases under realistic temperature regimes.

Daily temperature fluctuations significantly altered the thermal responses for all three adult mosquito traits studied here, primarily by reducing performance at temperatures near and above the thermal optimum. The reduced performance at warmer temperatures resulted in cooler upper thermal limits (*T_max_*) and thermal optima (*T_opt_*), and narrower thermal breadths, without a detectable impact on lower thermal limits (*T_min_*) (**Figure 2**, **Table 1**). Fluctuations also increased lifespan in our coldest mean temperature treatment (16°C). Our results contribute to a growing body of literature demonstrating that daily temperature fluctuations affect the life history of ectothermic organisms in ways not captured by constant mean temperature gradients^28,30–32,39,49,51,57–59^, including for mosquitoes and their associated pathogens^26,35–37,60–62^. The effect of temperature fluctuations on performance depends strongly on the mean temperature over which the fluctuation is occurring. Typically, fluctuations impair processes at the warmer end of the reaction norm and boost processes at the cooler end, resulting in cooler temperatures for both the *T_opt_* and *T_max_*, similar to our results. This general pattern is supported by three meta-analyses^28,57,58^ and frequently observed (albeit with some exceptions) in studies from medically important mosquitoes^35–37,61^ and other host-parasite systems^31,47^ (see **Table 4**). Collectively, these results suggest that whether fluctuations rescue or decrease performance is dependent on the mean temperature and the duration of time an organism remains beyond its thermal limits. Overall, our findings reinforce the pattern found in these previous studies: while fluctuations often reduce performance at warmer temperatures and increase it at cooler temperatures, there are also frequent exceptions to this rule.

**Table 4:**
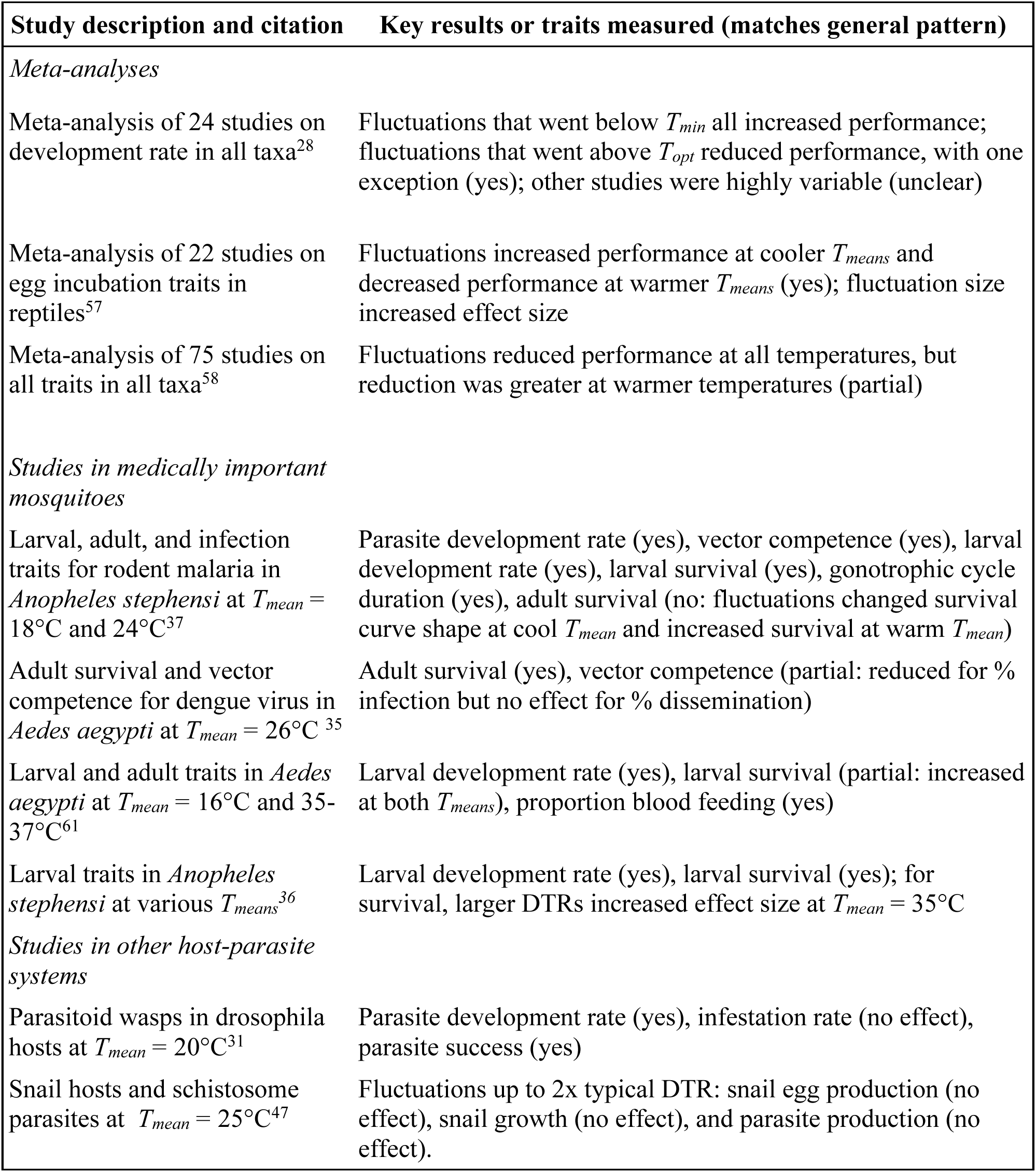
Summary of selected previous studies analyzing empirical data on the impact of temperature fluctuations on performance. When relevant, results include whether they match the general pattern of fluctuations improving performance at cooler temperatures and reducing performance at warmer temperatures. *T_mean_* = mean temperature, *T_min_* = lower thermal limit, *T_opt_* = thermal optimum, DTR = diurnal temperature range.

| Study description and citation | Key results or traits measured (matches general pattern) |
| --- | --- |
| *Meta-analyses* |  |
| Meta-analysis of 24 studies on development rate in all taxa<sup>28</sup> | Fluctuations that went below $T_{min}$ all increased performance; fluctuations that went above $T_{opt}$ reduced performance, with one exception (yes); other studies were highly variable (unclear) |
| Meta-analysis of 22 studies on egg incubation traits in reptiles<sup>57</sup> | Fluctuations increased performance at cooler $T_{means}$ and decreased performance at warmer $T_{means}$ (yes); fluctuation size increased effect size |
| Meta-analysis of 75 studies on all traits in all taxa<sup>58</sup> | Fluctuations reduced performance at all temperatures, but reduction was greater at warmer temperatures (partial) |
| *Studies in medically important mosquitoes* |  |
| Larval, adult, and infection traits for rodent malaria in *Anopheles stephensi* at $T_{mean} = 18^{\circ}\text{C}$ and $24^{\circ}\text{C}$<sup>37</sup> | Parasite development rate (yes), vector competence (yes), larval development rate (yes), larval survival (yes), gonotrophic cycle duration (yes), adult survival (no: fluctuations changed survival curve shape at cool $T_{mean}$ and increased survival at warm $T_{mean}$) |
| Adult survival and vector competence for dengue virus in *Aedes aegypti* at $T_{mean} = 26^{\circ}\text{C}$<sup>35</sup> | Adult survival (yes), vector competence (partial: reduced for % infection but no effect for % dissemination) |
| Larval and adult traits in *Aedes aegypti* at $T_{mean} = 16^{\circ}\text{C}$ and $35-37^{\circ}\text{C}$<sup>61</sup> | Larval development rate (yes), larval survival (partial: increased at both $T_{means}$), proportion blood feeding (yes) |
| Larval traits in *Anopheles stephensi* at various $T_{means}$<sup>36</sup> | Larval development rate (yes), larval survival (yes); for survival, larger DTRs increased effect size at $T_{mean} = 35^{\circ}\text{C}$ |
| *Studies in other host-parasite systems* |  |
| Parasitoid wasps in drosophila hosts at $T_{mean} = 20^{\circ}\text{C}$<sup>31</sup> | Parasite development rate (yes), infestation rate (no effect), parasite success (yes) |
| Snail hosts and schistosome parasites at $T_{mean} = 25^{\circ}\text{C}$<sup>47</sup> | Fluctuations up to 2x typical DTR: snail egg production (no effect), snail growth (no effect), and parasite production (no effect). |

Rate summation (RS) did not accurately predict trait values in diurnally fluctuating temperature environments in our study. Rate summation did predict reductions in performance near the thermal optima, but in many cases only captured a small proportion of the observed decrease (i.e. the direction of the effect was correct, but the magnitude was too small) (**Figure 3**, **Table 1**). Rate summation also predicted increases in performance near the thermal margins, yielding wider thermal breadths than what was observed, with both warmer *T_max_* and cooler *T_min_* values (i.e., the wrong direction of effect on *T_max_*). Finally, for our coldest mean temperature (16°C) and highest DTR (12°C), we observed lifespans that were higher than the maximum value observed for constant temperatures (i.e., at the *T_opt_*), something that is impossible to occur using rate-summation predictions. Few studies have quantitatively tested the predictions made by rate summation for how temperature fluctuation will alter organismal performance. One study using a green alga found that rate summation accurately predicted population growth rates in fluctuating conditions^30^. However, three studies on animals found that nonlinear averaging did not accurately predict performance of larval development and growth in frogs^39^, of short-term and long-term growth rates in tobacco hornworms^49^, and of development rate in coffin flies^28^. Alternatively, some studies compare their results qualitatively (i.e., did fluctuations increase or decrease performance) with general predictions based on Jensen’s Inequality and the concavity of the TPC. These studies typically find that the predicted change in trait value successfully matches the observed direction of trait change^51,63^ (but there are exceptions^59^). Thus, our findings once again reinforce general patterns from the literature: trait values measured under fluctuating conditions often qualitatively match the predicted changes compared to constant temperatures based on Jensen’s Inequality and the concavity of TPCs, but they rarely quantitatively match the specific values predicted by rate summation calculations.

Predicted thermal suitability for malaria transmission varied substantially among our four models (**Figure 4**, **Table 3**). These differences mirrored the trait-level results: empirical fluctuations (model 2) decreased *T_opt_* and *T_max_* compared to constant temperatures (model 1), while rate summation (models 4 and 5) predicted little change in *T_opt_*, increases in *T_max_*, and decreases in *T_min_*. Although rate summation appeared to predict the decrease in suitability at *T_opt_* quite accurately (**Table 3**), we note that our empirical model only accounts for the impact of fluctuations on our three focal traits, while the rate summation models simulate the impact of fluctuations on all seven traits. Thus, rate summation may still be only partially capturing the impact of fluctuating temperatures near the thermal optimum. We also found that the level at which rate summation was conducted had a large impact on predicted suitability near both thermal limits (i.e., on component traits versus on the TPC for suitability, model 4 versus model 5, respectively; **Figure 4C**). Studies on mosquito-borne disease generally perform rate summation on component traits^42–44,46^ or have ambiguously written methods^48^, but a recent study on temperature-dependent transmission of schistosomiasis performed rate summation directly on the TPC for *R_0_*^47^.

The variation in the suitability models’ thermal limits generated substantial differences in the predicted length of transmission seasons and geographic areas predicted to be suitable for year-round transmission (**Figures 5 and 6**, left columns). For our main mapping approach (number of months with *S(T)* > 0.001), constant temperatures (model 1) approximated empirically fluctuating temperatures (model 2) extremely well, since those models had the same thermal minima (*T_min_*). By contrast, both rate summation models (4 and 5) overpredicted the geographic area with year-round suitability for transmission due to their much cooler *T_min_*. The models’ upper thermal limits (*T_max_*) were not important here, since current monthly mean temperatures did not exceed them; however, they could begin to limit suitability under future climate change projections.

To our knowledge, this study is the first to attempt to validate the accuracy of rate summation in predicting the effects of thermal variation on mosquito and pathogen life history, and to explore the implications for predicted transmission. Unfortunately, our results suggest that these studies are likely overestimating transmission near *T_max_*, and possibly near *T_min_* as well. These areas of the TPC correspond to locations where the impacts of climate change on transmission are predicted to be felt most strongly, as cooler areas become newly suitable and hotter areas become unsuitable^3,7^. We found two studies that used rate summation to estimate the thermal response of transmission under multiple diurnally fluctuating conditions. Similar to our study, one predicted that fluctuations would increase the *T_max_* of transmission^48^, contrary to the decreases in *T_max_* observed in our study and that better correspond to the broader literature (**Table 4**). The other study provided results for a limited range of temperatures that stopped well below *T_max_* (at 28°C) and could not be compared^10^.

Overall, our suitability results are concerning: they demonstrate that rate summation calculations can systematically distort the thermal limits of TPCs and increase their thermal breadth, and yet many predictive models for mosquito-borne diseases use it to account for the impacts of temperature fluctuations on mosquito and pathogen traits that are important for transmission^10,35,42–46,48^. We recommend caution when applying rate summation to organismal performance and models for disease transmission (or other processes) in cases where empirical responses to fluctuating temperatures are not available. Rate summation more accurately estimates absolute levels of performance or transmission near the thermal optimum, which can be important for capturing the overall intensity of transmission. However, TPCs measured in constant temperature conditions may provide more accurate estimates of the thermal limits, which is important for estimating seasonality and the current and future geographic areas suitable for transmission. (This accuracy likely depends on the specific TPC function used: see below).

Many different factors could affect the accuracy of rate summation for predicting performance under fluctuating temperature conditions. First, the function chosen to fit the TPC over the constant temperature gradient will strongly influence any predictions from rate summation because the calculations are very sensitive to the shape and concavity of the thermal response, as illustrated by Jensen’s Inequality^27,40^. Many thermal responses are truncated at zero (including the quadratic and Brière responses used here) because negative values for traits like lifespan and fecundity are not biologically meaningful. This truncation, however, inherently creates accelerating (i.e., convex) portions of the curve, that in turn leads to higher predicted performance in fluctuating versus constant temperatures for mean temperatures near the thermal margins. TPCs that are not truncated below zero, such as the Eppley curve used in the study on population growth for a green alga, do not always predict an increase at the thermal margins using rate summation^30^. Second, traits that are rate-based (i.e., measured per unit time like development rate, foraging rate, daily fecundity, etc.) are more likely to show an asymmetrical thermal response^4^ and may be more likely to match the assumptions required for rate summation than traits that are integrated over an organism’s lifetime (e.g., longevity or lifetime fecundity). Third, traits that depend on discrete events may be determined by the temperatures an organism experiences shortly after those events occur. For example, the time of day mosquitoes are exposed to *Plasmodium falciparum* parasites and the portion of the DTR experienced after this exposure significantly alters the proportion of *Anopheles* mosquitoes that become infectious with malaria^60^. Finally, certain taxa may more closely match predictions from rate summation than others. For instance, rate summation may work better in single-celled organisms^30^ than in larger, multicellular organisms with more complex tissue-specific responses to temperature stress^32^. From a molecular and cellular biology perspective, discrepancies between observed performance and predictions from rate summation may occur due to acclimation/hardening processes or the accumulation of thermal stress and the energetic costs of repairing damage from extreme hot or cold temperatures^32,50,52,54^.

Organismal performance is consistently observed to differ in thermally fluctuating environments relative to constant temperature environments, thus developing a validated predictive framework that can accurately approximate trait performance in a fluctuating environment is essential. Future work should continue characterizing organismal responses in thermally fluctuating environments, in order to uncover potential patterns related to the type of trait and organism under study^26^. Additionally, we need more work that integrates phenomena across biological scales to mechanistically understand the cellular and molecular responses to thermal acclimation and stress that dictate the temperature constraints on organismal performance. Finally, while this study investigated the impact of thermal fluctuations on a single strain of mosquitoes in the adult stage, more work is needed to investigate how other environmental factors (e.g., food resources, competition, humidity) and genetic variation (e.g., thermal plasticity) affect organismal performance in thermally variable environments^6,50,64,65^.

In conclusion, realistic temperature fluctuations over the daily cycle can have significant impacts on organismal performance, including for mosquito vectors of human parasites like malaria. However, current approaches for quantitatively modeling the effect of temperature fluctuations using nonlinear averaging often fail to adequately predict performance under fluctuating conditions. Our thermal suitability model based on data from constant temperatures was more accurate for mapping the thermal limits for malaria transmission than the model parameterized via rate summation. Thus, for some applications it may be better to simply use thermal responses fit under constant temperature environments than to try to incorporate the impact of fluctuating temperatures using non-linear averaging. Future studies should carefully consider whether nonlinear averaging is likely to improve the accuracy of their results based on their specific goals. Meanwhile, more work is needed to improve methods for estimating the physiological effects of temperature fluctuation in real-world situations to more accurately predict organismal performance and disease transmission under realistic temperature regimes.

## Materials & Methods

### Mosquito husbandry

*Anopheles stephensi* mosquitoes (urban type form originally sourced from Walter Reed Army Institute of Research, Silver Spring, MD, USA) were reared at standard insectary conditions (27°C ± 0.5°C, 80% ± 5% relative humidity, and a 12L:12D photoperiod) prior to the life table experiment, as described previously ^8^. Briefly, we hatched immature mosquito larvae from eggs and placed 110 individuals into plastic trays (6 Qt., 12.4 cm x 34.6 cm x 21.0 cm) containing 500mL of distilled water. Food (100mg ground TetraMin fish flakes) was provided daily until most individuals reached the pupal stage. Pupae were rinsed and transferred to water-containing cups placed inside adult mosquito mesh cages for eclosion. For adult colony maintenance, *An. stephensi* were provided 5% dextrose and 0.05% para-amino benzoic acid (PABA) and fed whole human blood (O+, healthy male < 30 years, Interstate Blood Bank, TN, USA) via water-jacketed hog intestine membrane feeders to support reproduction.

### Experimental design

We adopted a similar experimental design as in ^8^, where we previously measured *An. stephensi* (urban type form) life history traits at six constant temperatures (16°C, 20°C, 24°C, 28°C, 32°C, and 36°C). Here, we programmed incubators (Percival; Perry, Iowa) to follow a Parton-Logan model ^66^ for hourly diurnal temperature ranges (DTR) that are relevant for *P. falciparum* transmission in a natural setting (DTR of 9°C or 12°C) around five of the mean temperatures (16°C, 20°C, 24°C, 28°C, 32°C ± 0.5°C) explored previously ^8^ (see **SI Methods**). All other incubator settings (80% ± 5 RH, and 12L:12D photoperiod) and experimental procedures were the same to allow for direct comparison between results. All experimental work for both studies was conducted during 2016-2018 at the University of Georgia (USA).

To generate a cohort of age-matched individuals, we collected pupae present at day nine post-hatch (when most immature mosquitoes reached the pupal stage) and placed them in an eclosion container within an adult cage for 24hr. We provided a sugar solution (5% dextrose and 0.05% para-amino benzoic acid) to co-housed age-matched adults for three days prior to starting the lifetable experiment to permit mating. The lifetable experiment was initiated by providing females with an initial blood meal for 15 min, randomly sorting 300 blood-fed females into individual housing (16oz. paper cup with mesh top), and then randomly assigning 30 individuals to each temperature treatment.

Each day until found dead, individuals were provided with a whole human blood meal for 15 minutes and inspected visually for imbibed blood. Oviposition sites (secured petri dish containing water saturated cotton and filter paper) within each individual housing were rehydrated and checked daily for eggs; if present, eggs were removed and counted. We terminated the experimental block when either all mosquitoes had died or when at most four mosquitoes remained alive at 16°C. The life table experiment for each fluctuation regime was performed two independent times resulting in data from a total of 600 individuals. Life table data collected across constant temperatures from the previous study by our group consisted of 390 individuals across six constant temperatures ^8^.

### Fitting Thermal Performance Curves (TPCs)

For each combination of trait (lifetime measures of bite rate [*a*], lifespan [*lf*], and egg production [*B*]) and fluctuation regime (constant, DTR 9°C, and DTR 12°C), we used a Bayesian framework to fit either a symmetric (quadratic: -*c*(*T-T_min_*)(*T-T_max_*)) or an asymmetric (Brière: *cT*(*T-T_min_*)(*T_max_*-*T*)^1/2^) non-linear unimodal function to generate a TPC predicting trait values across temperature (*T*, in degrees Celsius). From these functions, we can compare the predicted thermal limits (*T_min_*, *T_max_*) and optimum temperature (*T_opt_*) for each trait among the different DTR treatments, with *c* as a shape fit parameter. Both functions were restricted from becoming negative by assuming a trait value to be zero if *T* < *T_min_* or *T* > *T_max_.* The previous study^8^ analyzed only the constant temperature treatments and fit trait thermal responses to means from each experimental block using a truncated normal distribution. Here, we used the full dataset of three DTR treatments and fit the trait thermal responses to individual-level data, using different probability distributions for each trait based on the data type and observed distribution. For bite rate (*a*), we used a normal distribution truncated at zero; for lifespan (*lf*), we used a gamma distribution; for lifetime egg production (*B*), we used a negative binomial distribution (see **SI Methods** for model specifications).

For each trait, we selected the best-fitting functional form (quadratic or Brière) using the Deviance Information Criterion (DIC)^67^. For each parameter in the mean response function (i.e., *c*, *T_min_*, *T_max_*) and the additional parameter required to specify each probability distribution (i.e., the variance for the truncated normal distribution, the rate parameter for the gamma distribution, and the *r* parameter for the negative binomial distribution), we assumed low-information uniform priors (*T_min_* ∼ uniform (0, 20), *T_max_* ∼ uniform (28, 45), *c* ∼ uniform (0, 10), variance ∼ uniform (0,1000), rate ∼ uniform (1,100), r ∼ uniform (1,100)) that restricted the range of parameters to biologically or statistically meaningful values. TPCs were fitted in R using JAGS/R2jags^68,69^, which implements Markov Chain Monte Carlo (MCMC). Posterior draws were obtained from three concurrent Markov chains. In each chain, a 5,000-step burn-in phase was followed by 20,000 samples of the stationary chain, for a total of 60,000 posterior samples. These samples were thinned by saving every eighth sample (yielding 7,500 samples) to reduce autocorrelation in the chain. For each TPC, we used the posterior distributions for the parameters to generate posterior distributions over a temperature gradient from 0-45°C at 0.1°C intervals, which we then used to calculate the mean, median, and 95% credible intervals.

To test for the statistical significance of fluctuation treatment, we used the Deviance Information Criterion (DIC) output from JAGS. For each trait, we compared: 1) the sum of DIC values for the three models fit separately to data from each treatment (constant, DTR 9°C, and DTR 12°C) and 2) the DIC of a model fit to the combined data from all treatments. Fluctuation treatment is significant if the sum of the separate models is >= 2 DIC units lower than the DIC value for the combined model.

### Generating TPCs with rate summation

To calculate the trait thermal responses predicted by rate summation (**Equation 1**) we used the 7,500 posterior samples from the Bayesian fitted TPCs for each trait measured at constant temperatures. First, we used a Parton-Logan model^66^ to calculate a temperature profile for each mean temperature spanning 0-50°C with 0.1°C increments, assuming a DTR of 9 or 12°C across a 24-hour period (see **SI Methods**). Second, we calculated predicted trait values at each hour using the TPC for trait performance at constant temperatures. Third, a daily mean value for each trait was calculated by averaging the predicted hourly values for that trait over the 24-hour period for each mean temperature. When fluctuating temperatures extended beyond the range of our constant temperature TPCs (0°C ≥ T ≤ 45°C), we used the trait value predicted at the corresponding edge temperature, which was always equal or approximately equal to zero. Lastly, since rate summation was conducted for each posterior sample, we calculated the mean, median, and 95% credible interval of the resulting rate summation estimates for each mean temperature.

### Predicting thermal suitability, *S*(*T*)

Following previous work^8^, we use a modified expression for the relative pathogen basic reproductive number (relative *R_0_*), a metric of pathogen transmission potential in a given thermal environment. This metric incorporates the thermal responses of mosquito and parasite traits to evaluate the combined effects of temperature and temperature fluctuation on the predicted thermal suitability [*S(T)*, **Equation 2**] of *An. stephensi* to transmit *Plasmodium falciparum*^8^. A scaled version of *R_0_*(*T*), called *S(T)*, is proportional to the number of new cases expected to arise from a single case assuming a fully susceptible population, and is dependent on environmental temperature, *T* (°C). Further, because values for mosquito life history traits change as mosquitoes age, we have adopted the use of the *S(T)* expression that more precisely captures lifetime transmission potential^8^ (**Equation 2**).

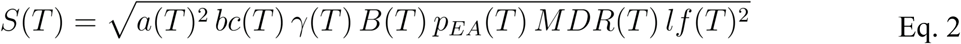

The parameters of *S(T)* include: daily per capita bite rate (*a*), vector competence (*bc*; the proportion of infectious mosquitoes), lifetime egg production (*B*), probability of egg-to-adult survival (*p_EA_*), mosquito development rate (*MDR*), and adult mosquito lifespan (*lf*). Further, the *S(T)* formulation uses the Gompertz function over daily adult survival and the extrinsic incubation period (*EIP*, the inverse of the parasite development rate [*PDR^−1^*]) to calculate the proportion of mosquitoes surviving the latency period (*ϒ*) as described in ^8^. We fit thermal responses for these additional traits (*p_EA_*, *MDR*, and *bc*) using previously published data measured across constant temperature gradients ^19,36^. For *ϒ*, we combined data for *PDR* measured across a constant temperature gradient ^19^ with our new lifespan (*lf*) data in constant and fluctuating conditions, and fit a TPC for each of our three fluctuation treatments (DTR = 0, 9, and 12°C). In all cases, we used the same methods as for the focal trait data collected here (described above), with a truncated normal distribution. We calculated thermal suitability using the full posterior distributions for each trait TPC over the temperature gradient from 0-45°C at 0.1°C intervals, yielding posteriors for suitability over that same gradient, with the same number of samples (7500). We then used these distributions to calculate the mean, median, and 95% credible intervals.

Absolute *R_0_(T)* is influenced by additional factors that we do not incorporate in this study including rainfall, humidity, mosquito habitat quantity and quality, infection status, and heterogeneity in contact rates, individuals, or genotypes. Thus, we instead describe the thermal suitability of pathogen transmission, *S(T)*, where *S(T)* is scaled to range between 0 and 1 at the respective minimum and maximum values for the median thermal response. We scaled all versions of the *S(T*) model using the maximum value from model version 1 (‘constant’, see *Suitability Model Overview*) in order to be able to visually compare differences in the predicted magnitude of thermal suitability between model versions. The additional *R_0_* parameters *r* (human recovery rate) and *N* (density of humans) are evaluated as arbitrary constants, as they are assumed to be temperature independent. Thus, a threshold of *S(T)* > 0 implies that the thermal conditions are suitable for the transmission of *P. falciparum* based solely on the temperature-dependent physiological responses of *An. stephensi*. Differences in the predicted critical temperatures at which *S(T)* reaches 0 (*T_min_* and *T_max_*) and 1 (*T_opt_*) can then be compared across diurnal temperature ranges.

### Sensitivity and uncertainty analysis

We performed two types of sensitivity analysis and an uncertainty analysis on each version of the suitability model to determine which traits were most important for determining the thermal optimum and limits for transmission and how each trait contributed to the uncertainty in *S(T)*. First, we used a partial derivative approach, calculating ∂*S*/∂*x*·∂*x*/∂*T* across the temperature (*T*) gradient for each trait (*x*). This approach only works for the models without rate summation (i.e., model 1: constant and model 2: empirical fluctuating) because it uses the derivatives of the quadratic and Brière functions and their fitted parameters (*T_min_*, *T_max_*, and *q*) for each trait. Second, we held each trait constant while allowing all others to vary with temperature. Finally, we calculated the HPD interval (highest posterior density interval, the smallest interval of predicted trait value encompassing 95% of the probability density in the posterior distribution) across the temperature gradient for *S*(*T*) using the full posterior distributions for all traits (i.e. full uncertainty) and for *S*(*T*) with each trait given its mean value (i.e. removing the uncertainty for one trait at a time). We then compared the relative size of the HPD in both conditions for each trait.

### Mapping thermal suitability predictions

We created maps to compare the spatial distribution of months of thermal suitability for transmission predicted by the different versions of our model, *S(T)*. For simplicity, we only mapped model versions 1, 2, 4, and 5 (constant, empirical fluctuating, trait-level RS fluctuating - all traits, and *S(T)*-level RS fluctuating, respectively) for one level of DTR (12°C) where applicable. As with previous mapping for thermal suitability of transmission ^5,8,23,42,70^, for each version of *S(T)* we determined the temperature range (at 0.1°C resolution) where *S(T)* > 0.001 with a posterior probability >97.5%. This conservative threshold minimizes type I error (inclusion of unsuitable areas). Here, we also calculated the temperature range at which each model exceeded an additional threshold of suitability, *S(T)* > 0.5. This threshold shows where thermal suitability is relatively high (rather than simply *present*), and allows us to illustrate quantitative differences between model versions 4 and 5 (i.e. rate summation performed on the trait TPCs versus on the suitability TPC), which had similar *T_min_* and *T_max_* but different shapes otherwise. For calculating the mapping thresholds, we scaled the 97.5% lower CI prediction from each model between 0 and 1 so that relative suitability was based on the maximum predicted suitability for that specific model.

Global gridded long-term average modeled baseline monthly mean temperatures at a 5 arcminute resolution (approximately 10 km2 at the equator), were downloaded from WorldClim.org (version 1.0). The number of months (0-12) of thermal suitability under each combination of model and suitability threshold was calculated at the pixel level, and masked to countries described as the ‘endemic’ range for *An. stephensi* (India, Pakistan, Iran, Kuwait, United Arab Emirates, and Oman), and for all countries in the continent of Africa, where it is currently invading and establishing. All raster calculations and mapping output were conducted in R (version 4.3.1), using packages ‘raster’ ‘terra’ ‘sf’ ‘tidyverse’ ‘ggplot2’ ‘maptools’ ‘mapdata’ ‘ggthemes’, in RStudio 2024.04.0 Build 735.

## Supporting information

Supplemental Methods and Results

## Data Availability

The mosquito trait data are currently available on the project GitHub repository: https://github.com/JoeyBernhardt/anopheles-rate-summation. Upon acceptance, these data will also be submitted to Dryad Data Repository, and the associated citation will be provided here.

## Code Availability

The code for this analysis is available on the project GitHub repository: https://github.com/JoeyBernhardt/anopheles-rate-summation.

## Acknowledgements

The authors would like to thank the VectorBite RCN Rate Summation Working Group and lab members for insightful discussion, as well as Francis Windram for technical assistance and code optimization. This research was supported by the NSF Graduate Research Fellowship Program (GRFP) and a NIH R01 award (1R01AI110793-01A1). MSS was supported by NSF EID: Effects of Temperature on Vector-borne Disease Transmission (DEB-1518681). SJR was supported by NSF CIBR: VectorByte: A Global Informatics Platform for studying the Ecology of Vector-Borne Diseases (NSF DBI 2016265).

## Author Contributions

KLM and CCM designed the study, with input from RJH. KLM and ARO performed the experiments. KLM performed the first analysis and wrote the first manuscript. MSS and JRB revised the analysis, with input from CCM and VMS. SJR performed the mapping analysis. MSS and CCM revised the manuscript, with input from SJR and VMS. All authors read and approved the final manuscript.

## Notes

https://github.com/JoeyBernhardt/anopheles-rate-summation

### Competing Interest Statement

The authors have declared no competing interest.

