## Supplemental Methods and Results for "Mean daily temperatures can predict the thermal limits of malaria transmission better than rate summation"

#### Table of Contents

1. **Table S1**: Deviance information criterion (DIC) values used to compare TPCs for adult mosquito traits fit with quadratic and Brière functions.
2. **Table S2**: Deviance information criterion (DIC) values used to compare fluctuation treatments for adult mosquito traits.
3. **Table S3**: Properties of thermal performance curves (TPCs) for adult mosquito traits in constant and fluctuating temperatures
4. **Table S4**: Predicted thermal suitability for transmission for five different models.
5. **Table S5**: Properties of thermal performance curves (TPCs) for other mosquito and pathogen traits using data from previous studies in constant conditions.
6. **Figure S1**: Sensitivity Analysis 1 of suitability models – partial derivatives.
7. **Figure S2**: Sensitivity Analysis 2 of suitability models – holding single parameters constant.
8. **Figure S3**: Uncertainty Analysis of suitability models.
9. **Supplemental Methods**:
    a. Trait TPC model specifications
    b. Parton-Logan model for diurnal temperature fluctuations

**Table S1. Deviance information criterion (DIC) values used to compare TPCs for adult mosquito traits fit with quadratic and Brière functions.** The lower of the two DIC values (quadratic or Brière model) and ΔDIC >2 are bolded.

| Trait & Fluctuation Regime | Quadratic model | Brière model | ΔDIC |
|---|---|---|---|
| Bite rate (*a*) | | | |
| *Constant* | -449.7 | **-492.4** | **42.7** |
| *DTR 9* | -462.3 | -471.5 | **9.2** |
| *DTR 12* | -435.2 | -445.1 | **9.9** |
| Lifespan (*lf*) | | | |
| *Constant* | **3150.9** | 3339.9 | **189.0** |
| *DTR 9* | **2436.5** | 2533.1 | **96.5** |
| *DTR 12* | 2521.5 | 2533.1 | **11.6** |
| Lifetime eggs (*B*) | | | |
| *Constant* | **4586.7** | 4587.8 | 1.1 |
| *DTR 9* | 3507.9 | **3506.4** | 1.5 |
| *DTR 12* | **3294.5** | 3296.4 | 1.9 |

**Table S2. Deviance information criterion (DIC) values used to compare fluctuation treatments for adult mosquito traits.** Models were fit to data from each fluctuation treatment separately, data from all treatments combined ("Combined model"), and data for both fluctuating treatments combined ("DTR combined model"). Statistically significant DIC values are bolded. Fluctuation treatment was statistically significant for all traits. The magnitude of the fluctuation was only significant for lifespan. Medians

| Trait | Constant model | DTR9 model | DTR12 model | Sum of all separate models | Combined model | Sum of DTR models | DTR combined model |
|---|---|---|---|---|---|---|---|
| Bite rate (*a*) | -496.4 | -476.4 | -449.4 | **-1422.1** | -1340.9 | -925.7 | -925.5 |
| Lifespan (*lf*) | 3146.2 | 2431.4 | 2516.3 | **8094.0** | 8166.2 | **4947.8** | 4953.4 |
| Lifetime eggs (*B*) | 4581.1 | 3503.0 | 3289.7 | **11373.8** | 11385.2 | 6792.7 | 6793.1 |
| Gamma (Ɣ) | -28.9 | -29.0 | -21.3 | **-79.1** | -68.2 | -50.3 | -48.9 |

**Table S3. Properties of thermal performance curves (TPCs) for adult mosquito traits in constant and fluctuating temperatures.** For directly fitted TPCs, parameters ($q$, $T_{min}$ = thermal minimum, and $T_{max}$ = thermal maximum) are for Brière (bite rate) or quadratic (lifespan and lifetime egg production) functions. The remaining TPCs were calculated via rate summation (RS). Diurnal temperature ranges (DTR) = 9 and 12°C. Values are medians of MCMC posteriors (95% credible intervals in parentheses). See main text Figure 2.

| **Trait & Fluctuation Treatment** | ***q*** **(°C)** | $T_{min}$ **(°C)** | $T_{max}$ **(°C)** | $T_{opt}$ **(°C)** | $T_{breadth}$ **(°C)** |
|---|---|---|---|---|---|
| Bite rate (*a*) | | | | | |
| *Constant* | $1.62 \cdot 10^{-4}$ ($1.39$–$2.06 \cdot 10^{-4}$) | 2.30 (0.11–6.26) | 42.2 (40.5–44.3) | 34.0 (32.8–35.6) | 40.0 (34.6–43.6) |
| *Empirically fit DTR 9* | $1.66 \cdot 10^{-4}$ ($1.41$–$2.13 \cdot 10^{-4}$) | 1.55 (0.06–5.38) | 37.0 (35.5–39.3) | 29.8 (28.7–31.5) | 35.5 (30.5–38.6) |
| *Empirically fit DTR 12* | $1.52 \cdot 10^{-4}$ ($1.19$–$2.15 \cdot 10^{-4}$) | 2.66 (0.13–7.4) | 39.1 (36.5–43.4) | 31.6 (29.8–34.9) | 36.6 (29.5–42.3) |
| *RS DTR 9* | NA | 0.0 (0.0–1.2) | 45.0 (44.5–45.0) | 33.0 (31.8–34.7) | 45.0 (43.5–45.0) |
| *RS DTR 12* | NA | 0.0 (0.0–0.0) | 45.0 (45.0–45.0) | 32.2 (31.0–34.0) | 45.0 (45.0–45.0) |
| Lifespan (*lf*) | | | | | |
| *Constant* | 0.10 (0.09–0.12) | 1.04 (0.04–3.82) | 38.9 (38.3–39.8) | 20.0 (19.4–21.3) | 37.9 (34.8–39.4) |
| *Empirically fit DTR 9* | 0.12 (0.10–0.15) | 1.09 (0.48–4.24) | 35.8 (34.9–36.9) | 18.5 (17.8–19.9) | 34.7 (31.1–36.49) |
| *Empirically fit DTR 12* | 0.14 (0.12–0.16) | 0.73 (0.02–3.12) | 35.0 (34.2–35.9) | 17.9 (17.3–19.0) | 34.2 (31.6–35.5) |
| *RS DTR 9* | NA | 0.0 (0.0–0.0) | 43.0 (42.3–43.8) | 20.0 (19.4–21.3) | 43.0 (42.3–43.8) |
| *RS DTR 12* | NA | 0.0 (0.0–0.0) | 44.3 (43.6–45.0) | 20.1 (19.4–21.3) | 44.3 (43.6–45.0) |
| Lifetime eggs (*B*) | | | | | |
| *Constant* | 2.08 (1.13–3.13) | 12.4 (6.87–14.1) | 37.7 (36.9–39.8) | 25.0 (22.8–25.9) | 25.4 (23.0–32.3) |
| *Empirically fit DTR 9* | 2.33 (0.81–4.30) | 12.3 (4.68–14.3) | 35.2 (33.4–42.5) | 23.8 (21.4–26.0) | 23.1 (19.4–35.7) |
| *Empirically fit DTR 12* | 2.16 (0.72–4.26) | 11.9 (2.29–14.5) | 34.8 (33.1–41.1) | 23.4 (19.7–25.1) | 23.0 (19.0–36.6) |
| *RS DTR 9* | NA | 7.3 (1.8–9.1) | 41.7 (40.9–43.8) | 25.1 (22.8–26.0) | 32.1 (34.5–41.3) |
| *RS DTR 12* | NA | 5.7 (0.2–7.5) | 43.0 (42.2–45.0) | 25.1 (22.8–26.0) | 37.4 (35.0–43.6) |

**Table S4. Predicted thermal suitability for transmission for five different models.** Properties of the thermal performance curves: thermal minimum ($T_{min}$), thermal maximum ($T_{max}$), thermal optimum ($T_{opt}$), and thermal breadth ($T_{breadth}$). Fluctuating models are parameterized with trait TPCs fit directly from empirical data ("Empirical fluctuating") or are calculated using rate summation (RS). Rate summation was used only for the three traits with empirical data ("Trait-level RS - 3 traits"), for all traits ("Trait-level RS - all traits"), or directly on the TPC for suitability at constant temperatures ("*S*(*T*)-level RS"). Diurnal temperature ranges (DTR) = 9 and 12°C. Values are medians of posteriors (95% credible intervals in parentheses). See main text Figure 4.

| **Suitability Model** | $T_{min}$ **(°C)** | $T_{max}$ **(°C)** | $T_{opt}$ **(°C)** | $T_{breadth}$ **(°C)** |
|---|---|---|---|---|
| Constant | 15.0 (14.3–15.7) | 36.0 (35.1–36.0) | 26.8 (26.5–27.2) | 21.0 (20.2–21.7) |
| Empirical fluctuating | | | | |
| *DTR 9* | 15.0 (14.3–15.7) | 35.2 (33.4–36.1) | 25.6 (25.2–26.1) | 20.2 (18.2–21.5) |
| *DTR 12* | 15.0 (14.3–15.7) | 34.8 (33.2–36.1) | 25.4 (25.0–25.9) | 19.8 (17.9–21.5) |
| Trait-level RS - 3 traits | | | | |
| *DTR 9* | 15.0 (14.3–15.7) | 36.0 (35.9–36.1) | 26.7 (26.4–27.1) | 21.0 (20.2–21.7) |
| *DTR 12* | 15.0 (14.3–15.7) | 36.0 (35.9–36.1) | 26.6 (26.3–27.0) | 21.0 (20.2–21.7) |
| Trait-level RS - all traits | | | | |
| *DTR 9* | 10.0 (9.3–10.7) | 40.0 (39.9–40.1) | 26.6 (26.2–26.9) | 30.0 (29.2–30.7) |
| *DTR 12* | 8.4 (7.7–9.1) | 41.3 (41.2–41.4) | 26.4 (26.0–26.7) | 32.9 (32.1–33.6) |
| *S*(*T*)-level RS | | | | |
| *DTR 9* | 10.0 (9.3–10.7) | 40.0 (39.9–40.1) | 26.7 (26.4–27.1) | 30.0 (29.2–30.7) |
| *DTR 12* | 8.4 (7.7–9.1) | 41.3 (41.2–41.4) | 26.9 (26.6–27.4) | 32.9 (32.1–33.6) |

**Table S5. Properties of thermal performance curves (TPCs) for other mosquito and pathogen traits using data from previous studies in constant conditions.** Parameters ($q$, $T_{min}$ = thermal minimum, and $T_{max}$ = thermal maximum) are for Brière (*PDR* and *MDR*) or quadratic (vector competence and egg-to-adult survival) functions. Values are medians of MCMC posteriors (95% credible intervals in parentheses).

| **Trait & data source** | **$q$ (°C)** | **$T_{min}$ (°C)** | **$T_{max}$ (°C)** | **$T_{opt}$ (°C)** | **$T_{breadth}$ (°C)** |
|---|---|---|---|---|---|
| Pathogen dev. rate (*PDR*)[19] | $5.08 \cdot 10^{-5}$ ($4.10$–$6.89 \cdot 10^{-5}$) | 8.59 (4.23–12.3) | 43.9 (40.7–45.0) | 36.0 (33.8–36.9) | 35.3 (29.0–40.1) |
| Vector competence (*bc*)[19] | $2.22 \cdot 10^{-3}$ ($1.24$–$5.15 \cdot 10^{-3}$) | 8.00 (0.57–15.4) | 40.0 (36.6–44.3) | 24.2 (20.8–26.9) | 32.2 (21.8–42.2) |
| Prob. egg to adult survival ($p_{EA}$)[36] | $7.51 \cdot 10^{-3}$ ($6.45$–$8.53 \cdot 10^{-3}$) | 15.1 (14.3–15.7) | 37.3 (36.7–38.1) | 26.2 (25.8–26.6) | 22.3 (21.3–23.5) |
| Mosquito dev. rate (*MDR*)[36] | $1.06 \cdot 10^{-4}$ ($1.00$–$1.13 \cdot 10^{-4}$) | 13.3 (12.6–14.0) | 36.0 (35.9–36.0) | 30.5 (30.4–30.6) | 22.6 (22.0–23.4) |

**Figure S1: Sensitivity Analysis 1 of suitability models – partial derivatives.** This approach only works for the models without rate summation (i.e., model 1: Constant T and model 2: Empirical Fluctuating T) because it uses the derivatives of the quadratic and Brière functions and the fitted parameters (Tmin, Tmax, and q) for each trait. See *Methods* for details. (A) Model 1: constant temperature, (B) model 2: empirical DTR 9, (C) model 2: empirical DTR 12.

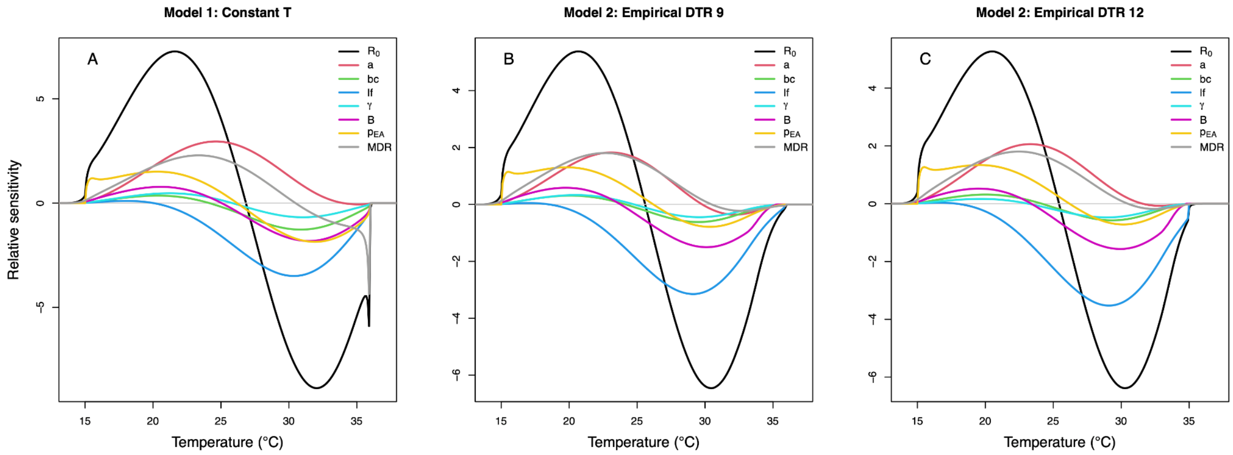

**Figure S2: Sensitivity Analysis 2 of suitability models – holding single parameters constant.** (A) Model 1: constant temperature, (B) model 2: empirical DTR 9, (C) model 2: empirical DTR 12, (D) model 3: trait-level rate summation - 3 traits for DTR 9, (E) model 3: trait-level rate summation - 3 traits for DTR 12, (F) model 4: trait-level rate summation - all traits for DTR 9, (G) model 4: trait-level rate summation - all traits for DTR 12, (H) model 5: S(T)-level rate summation for DTR 9, and (I) model 5: S(T)-level rate summation for DTR 12.

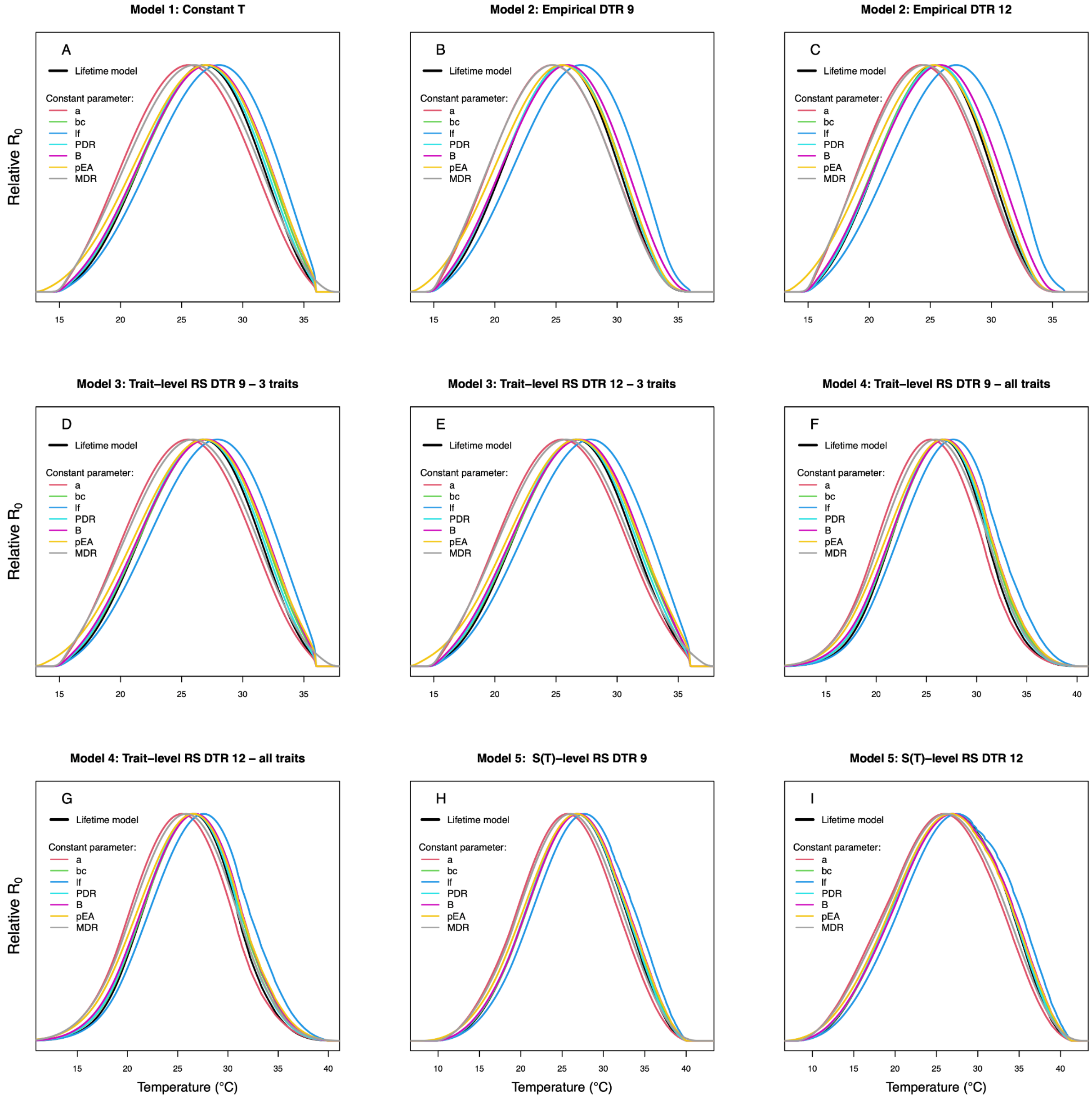

**Figure S3: Uncertainty Analysis of suitability models.** The width in credible intervals due to each parameter. (A) Model 1: constant temperature, (B) model 2: empirical DTR 9, (C) model 2: empirical DTR 12, (D) model 3: trait-level rate summation - 3 traits for DTR 9, (E) model 3: trait-level rate summation - 3 traits for DTR 12, (F) model 4: trait-level rate summation - all traits for DTR 9, (G) model 4: trait-level rate summation - all traits for DTR 12, (H) model 5: S(T)-level rate summation for DTR 9, and (I) model 5: S(T)-level rate summation for DTR 12.

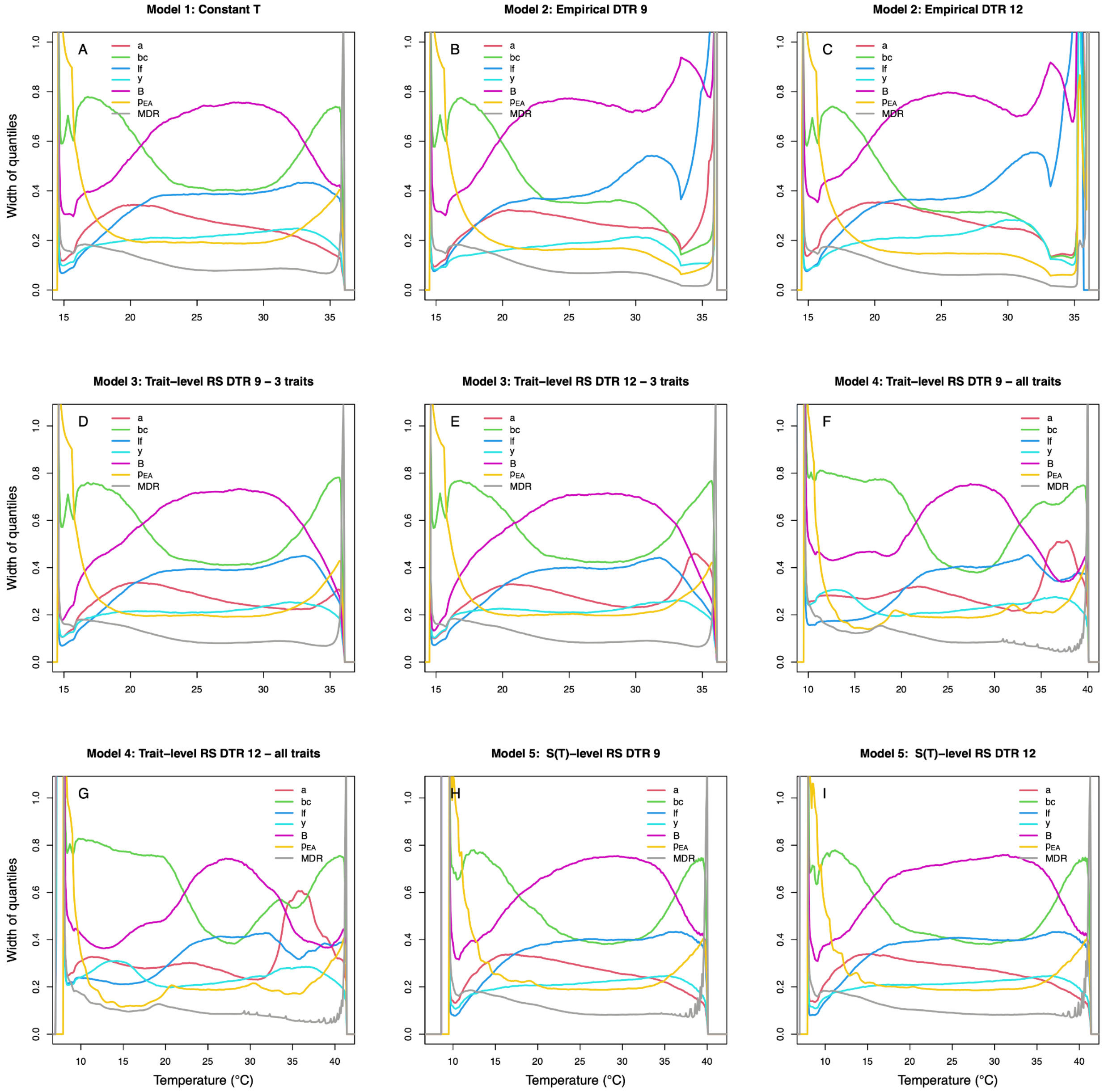

### Supplemental Methods

#### Trait TPC model specifications

In the following models, the subscript *i* denotes values corresponding to each individual-level observation. The temperature-dependent mean trait value ($\mu_i$) is defined by either a quadratic or Brière function. The inequalities are used to restrict the trait values to zero where the temperature is greater than $T_{max}$ or less than $T_{min}$. The other parameters (τ, s, r, and p) are used to define the relevant probability distribution and relate it to $\mu_i$. The corresponding code can be found in the project GitHub repository.

*Normal likelihood truncated at 0 with Brière function for bite rate*

bite $\text{rate}_i \sim \text{normal}(\mu_i , \tau)$ truncated(0,)

$\mu_i = cT(T\text{-}T_{min})(T_{max}\text{-}T)^{1/2})(T > T_{min})(T < T_{max})$

*Negative binomial likelihood with quadratic function for lifespan*

$\text{lifespan}_i \sim \text{gamma}(s_i, r)$

$s_i = r * \mu_i$

$\mu_i = -c(T\text{-}T_{min})(T\text{-}T_{max}))(T > T_{min})(T < T_{max})$

*Gamma likelihood with quadratic function for lifetime egg production*

$\text{egg}_i \sim \text{negative binomial}(p_i, r)$

$p_i = r / (r + \mu_i )$

$\mu_i = -c(T\text{-}T_{min})(T\text{-}T_{max}))(T > T_{min})(T < T_{max})$

**Parton-Logan model for diurnal temperature fluctuations**

To simulate a stereotypical profile of temperature fluctuation around a temperature mean through the course of a 24-hour period, we programmed our incubator using a Parton-Logan model [66] to run 10 daily temperature fluctuation profiles. Profiles fluctuated at the hourly scale with a diurnal temperature range (DTR) of either 9 or 12°C around five mean temperatures (16, 20, 24, 28, or 32°C; **Figure 1**). Parton-Logan models assume a sinusoidal relationship with temperature during the day ($T_{day}$) that peaks at a maximum temperature ($T_{max}$), followed by an exponential decay during the night ($T_{night}$; **Equation S1**) that asymptotes at the minimum temperature ($T_{min}$).

$$T_{day}(m) = T_{min} + (T_{max} - T_{min}) \sin\left(\frac{\pi m}{Y + 2a}\right) \quad \text{Eq. S1A}$$

$$T_{night}(n) = \frac{T_{min} - T_{sunset} \cdot e^{\frac{-Z}{\tau}} + (T_{sunset} - T_{min}) e^{\frac{-n}{\tau}}}{1 - e^{\frac{-Z}{\tau}}} \quad \text{Eq. S1B}$$

Note these definitions of $T_{min}$ and $T_{max}$ differ from those used elsewhere in the paper, which describe the thermal limits of a TPC. For a given temperature regime, $T_{sunset}$ is calculated as the final temperature predicted by **Equation S1A** and then used to parameterize **Equation S1B**. We assumed a day length ($Y$) and night length ($Z$) of 12 hours each, such that the ratio of light to dark hours was 12:12. The day constant ($a$ = 1.5) sets the period of the sine wave and thus determines the timing of $T_{max}$, (where larger $a$ delays $T_{max}$), while the night constant ($\tau$ = 4) adjusts the timing of the exponential decay (where smaller $\tau$ results in a faster decay). The inputs $m$ and $n$, denote the number of hours after sunrise and sunset, respectively, are used to calculate the desired sequence of hourly temperatures over time. Finally, the mean of *Tmin* and *Tmax* do not exactly equal the mean daily temperature predicted by this model.

With this model, the mean daily temperature ($T_{mean}$) is close but not exactly equal to the mean of $T_{min}$ and $T_{max}$. Thus, in order to parameterize the model for a specific mean temperature, you cannot determine $T_{min}$ and $T_{max}$ simply by subtracting or adding (respectively) half the DTR to the desired $T_{mean}$ – a small corrective factor ($c$) must also be used. This corrective factor is scaled by the DTR and is specific to a given day length. For a 12:12 day:night cycle, $c$ = 0.0575824 and is used according to:

$$T_{min} = T_{mean} - \frac{DTR}{2} - c \cdot DTR \qquad \text{Eq. S2A}$$

$$T_{max} = T_{mean} + \frac{DTR}{2} - c \cdot DTR \qquad \text{Eq. S2B}$$
